# Oxaloacetate damages mitochondria by perturbing MIC60-dependent membrane remodeling

**DOI:** 10.64898/2026.01.28.702225

**Authors:** Jie Zhang, Qian Shan, Xin Wang, Meijiao Li, Yang Yang, Mei Duan, Ruofeng Tang, Junxiang Zhou, Fengyang Wang, Yuehui Shi, Kai Jiang, Chonglin Yang

## Abstract

Mitochondria catabolize nutrients by generating sequentially-ordered organic acid intermediates that are oxidized through the tricarboxylic acid cycle. Pathogenic accumulation of metabolic organic acids manifests as devastating organic acidemias/acidurias and other severe diseases, but the underlying mechanisms are largely unknown. Using unbiased *C. elegans* genetic screening, we here reveal that mutations in the phosphoenolpyruvate carboxykinases PCK-1 and PCK-2 cause buildup of oxaloacetate, a key tricarboxylic acid cycle intermediate, leading to severe mitochondrial damage. Depletion of mitochondrial GOT-2.1 or GOT-2.2, which catalyze oxaloacetate conversion to aspartate, also causes oxaloacetate accumulation and defective mitochondria with disrupted cristae. We demonstrate that oxaloacetate binds the MICOS complex subunit CHCH-3/MIC19 and inhibits its function of promoting IMMT-1/MIC60-dependent membrane shaping and remodeling. In mammalian cells, aberrant OAA buildup similarly causes mitochondrial impairment through MIC19 and MIC60. These findings not only provide important mechanistic insights into mitochondrial damage in the context of defective oxaloacetate metabolism, but also suggest therapeutic strategies for oxaloacetate-related mitochondriopathies.

## Introduction

Mitochondria are double-membrane organelles that are fundamental to cellular metabolism. On one hand, mitochondria oxidize nutrients, including glucose, amino acids and fatty acids, through the tricarboxylic acid (TCA) cycle, which is coupled with oxidative phosphorylation to generate ATP, providing energy for various cellular activities[1]. On the other hand, mitochondria synthesize diverse biomolecules and are indispensable for maintenance of cellular ion homeostasis[1]. To carry out such complex metabolic activities, mitochondria are highly compartmentalized by cristae, which are formed by inward invagination of the inner mitochondrial membranes[2]. The formation and maintenance of the dynamic cristae structure require coordinated actions of many factors, including the MICOS complex, respiratory chain complexes and the ATP synthase complex[3, 4]. Perturbing the functions of these major factors causes damage to mitochondrial architecture, leading to functional impairment of mitochondria. Mitochondrial dysfunction is involved in many human diseases, including inborn errors of metabolism, neurodegenerative diseases, cardiomyopathies, diabetes and aging[5].

At the center of mitochondrial metabolism, the TCA cycle is sustained by multiple intermediate organic acids. In addition to their key role in cellular energy production, these acid intermediates serve as precursors for biosynthesis and participate in protein post-translational modifications and epigenetic regulation of gene expression[6]. Insufficiency or excess of these intermediates not only affects the TCA cycle, but also causes changes of mitochondrial architecture[7–9]. In fact, aberrant accumulation of mitochondrial organic acids often leads to devastating organic acidemias/acidurias, with symptoms including global developmental delay and early onset of neurologic problems[10]. Among the TCA cycle intermediates, OAA is particularly important because it acts as both the start and end points of each TCA cycle. OAA is condensed with acetyl-CoA to generate citrate for initiation of the TCA cycle, and it is regenerated from malate (Mal) by the malate dehydrogenase 2 (MDH2) at the end[11]. In cases where TCA intermediates are insufficient, OAA can be replenished from pyruvate by the enzyme pyruvate carboxylase[12, 13]. Because the TCA cycle only uses a small proportion of OAA[14], a significant proportion of mitochondrial OAA is converted to aspartate (Asp) by the mitochondrial glutamic-oxaloacetic transaminase 2 (GOT2), and is then transported out of mitochondria through the aspartate-malate shuttling system[15–17]. In the cytosol, OAA is generated by multiple mechanisms, including conversion from Mal by the malate dehydrogenase 1 (MDH1), conversion from Asp by the GOT1 transaminase, and generation from citrate by the ATP-citrate lyase (ACLY)[18–20]. Cytosolic OAA can be converted to Asp to participate in the urea cycle, nucleotide synthesis and amino acid synthesis[21]. More importantly, OAA serves as the precursor of gluconeogenesis, in which it is converted to phosphoenolpyruvate (PEP) by phosphoenolpyruvate carboxykinase (PEPCK)[22–24]. Because OAA lies at the nexus of catabolism and anabolism, disruption of OAA metabolism is closely linked to human disorders. For example, deficiency in pyruvate carboxylase causes multiorgan metabolic imbalance, which manifests as lactic academia and neurological dysfunction at an early age[25–27]. Mutations of MDH2 cause early-onset severe encephalopathy[28–30]. Patients with GOT2 mutations display global developmental delay, epilepsy, and progressive microcephaly[15, 31]. In addition, mutations of the cytosolic PEPCK, encoded by the *PCK1* gene, lead to defective gluconeogenesis and hypoglycemia, liver failure, and developmental delay[32–34]. Nevertheless, it is not fully understood whether and how disruption of OAA metabolism adversely affects mitochondrial architecture and function to contribute to the pathogenesis of OAA metabolism-related human disorders.

Given that many metabolic pathways, including the TCA cycle, are evolutionarily conserved across diverse species, we have established the genetically tractable *Caenorhabditis elegans* as an outstanding model to explore how metabolic intermediates adversely affect mitochondrial structure and functions[7, 8, 10]. Here, combining unbiased genetic screening and targeted gene depletion in *C. elegans*, we uncover that aberrant accumulation of cytosolic and mitochondrial OAA leads to structural and functional mitochondrial defects. Mechanistically, we demonstrate that OAA acts through the CHCH-3/MIC19 subunit of the MICOS complex to cause mitochondrial damage. Moreover, we show that aberrantly accumulated OAA impairs mitochondria in mammalian cells in a MICOS-dependent manner similar to that in *C. elegans*. These findings thus provide important novel insights into maintenance of mitochondrial homeostasis and human pathologies related to OAA metabolism.

## Results

### Loss of *pck-2* causes mitochondrial abnormalities

By mutagenizing *C. elegans* hermaphrodites expressing MITO-GFP and screening mutants with abnormal mitochondrial morphology, we identified two mutants, *yq270* and *yq271*, together with many other mutants that displayed varying abnormal mitochondrial morphologies compared to the predominantly tubular mitochondria in the N2 wild-type animals[7, 8, 35]. *yq270* and *yq271* mutants were characterized by the appearance of enlarged and spherical mitochondria in hypodermal cells in a development-dependent manner (S1A Fig). The mitochondrial defect was confirmed by co-labeling mitochondria with MITO-GFP and mCherry-tagged TOMM-20 (TOMM-20::mCh), an outer mitochondrial membrane protein (Fig 1A). Spherical mitochondria also existed in the body-wall muscle cells and intestinal cells in these mutants (Fig 1B). *yq270* and *yq271* animals showed significant reduction in ATP levels and body-bending rates (Fig 1C and 1D). In addition, both mutants exhibited a lower percentage of hatched embryos, reduced lifespan and shorter body lengths (Fig 1E-1G). Together these results suggest that the *yq270* an *yq271* mutations cause mitochondrial abnormalities, leading to defective development and growth.

**Fig 1.**
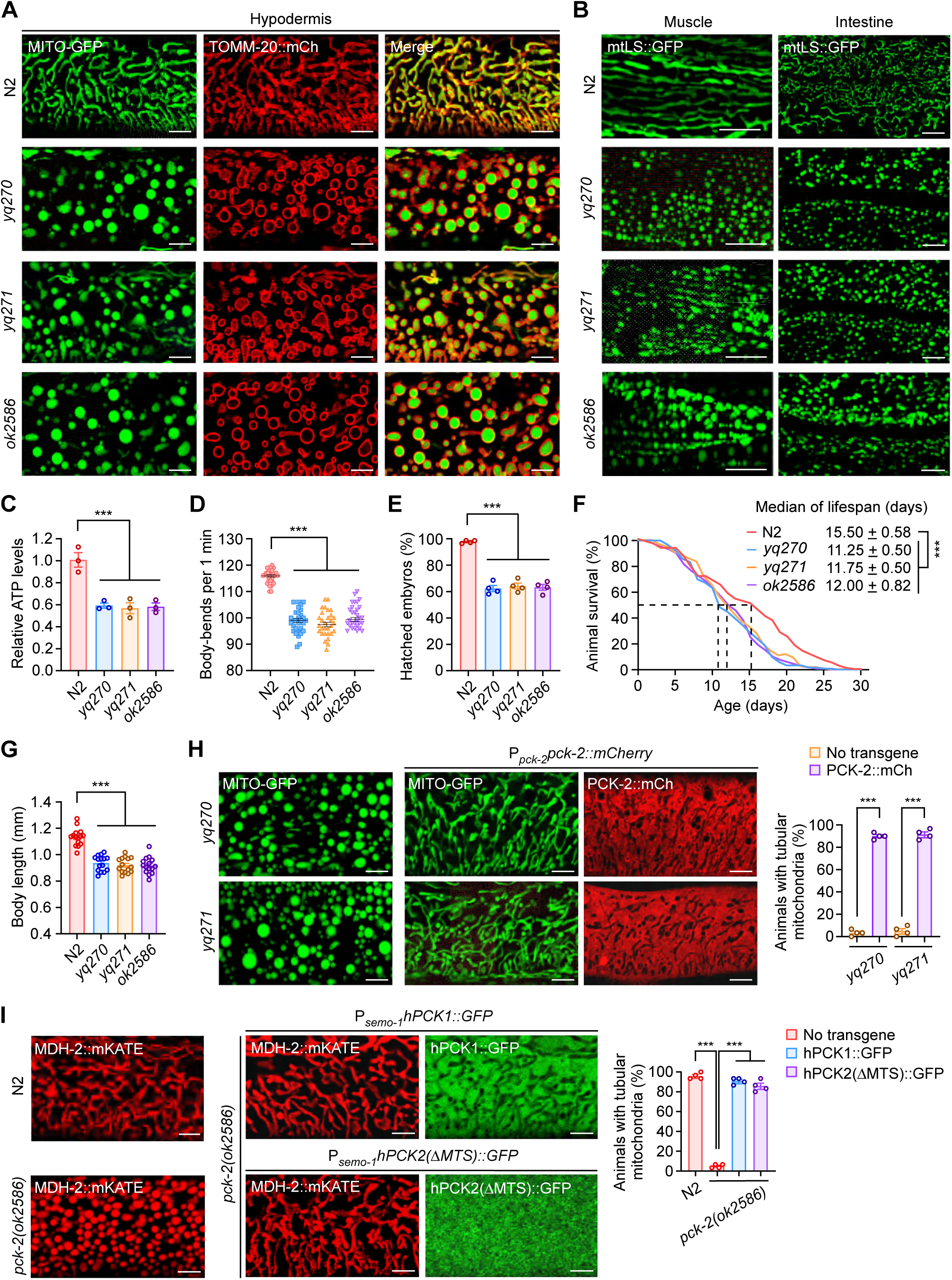
*pck-2* mutations cause abnormal mitochondrial morphology and functions. **(A)** Representative images of mitochondria co-labeled with MITO-GFP and TOMM-20::mCh (mCherry) in the hypodermis in N2 (wild type) and *yq270*, *yq271* and *ok2586* mutants at the adult stage (48 h after the 4^th^ larval (L4) stage). Bars, 5 μm. **(B)** Representative images of mitochondria labeled with MITO-GFP in the muscle and intestine in N2 and *yq270*, *yq271* and *ok2586* mutants at 48 h post L4 stage. Bars, 5 μm. **(C-G)** Relative ATP levels (300 animals were analyzed for each genotype, *n*= 3. independent experiments) (C), body-bending rates (*n*=30 animals were analyzed for each genotype) (D), percentage of hatched embryos (4 animals were analyzed for each genotype, *n*= 4 independent experiments) (E), adult lifespan (120 animals were analyzed for each genotype) (F) and adult body length (*n*=15 animals were analyzed for each genotype) (G). **(H)** Images (left) and quantification (right) of the rescue of abnormal mitochondria (labeled with MITO-GFP) in the hypodermis in *pck-2(yq270)* and *pck-2(yq271)* mutants by transgenic PCK-2::mCh driven by the *pck-2* promoter (P*_pck-2_pck-2::mCh*). A total of 120 animals were examined for each genotype in *n*= 4 independent experiments. Bars, 5 μm. **(I)** Images (left and middle) and quantification (right) of the rescue of abnormal mitochondria (labeled with MDH-2::mKATE) in the hypodermis in *pck-2(ok2586)* mitochondria by transgenic hPCK1::GFP (P*_semo-1_PCK1::gfp*) or hPCK2(ΔMTS)::GFP (P*_semo-1_(ΔMTS)PCK2::gfp*) driven by the *semo-1* promoter. A total of 120 animals were examined for each genotype in *n*= 4 independent experiments. Bars, 5 μm. For quantifications, data points represent (mean ± s.e.m.). *P* values were determined using one-way ANOVA in (C-G), two-tailed unpaired Student’s *t*-test in (H) or two-way ANOVA in (I). \*\*\**P*<0.001; \*\**P* < 0.01; \**P* < 0.05; ns, not significant (*P* > 0.05).

Genetic mapping and sequencing revealed that *yq270* and *yq271* both cause mutations in the *pck-2* gene on linkage group I (LGI). *yq270* (g.4060 G>A) results in a premature stop codon at W212 of the PCK-2 protein, while *yq271* (g.4375 C>T) results in a S317F missense mutation (S1B and S1C Fig). *C. elegans* PCK-2 and its homolog PCK-1 are homologous to the mammalian PEPCKs PCK1 and PCK2 (S1C Fig), which catalyze the conversion of OAA to PEP in gluconeogenesis[24]. Animals carrying a previously identified *pck-2* deletion allele, *pck-2(ok2586)*, similarly exhibited spherical mitochondria, impaired mitochondrial functions, and defective development and growth (Fig 1A-1G). We created a rescue construct consisting of transgenic PCK-2 tagged with mCherry, with expression driven by the *pck-2* promoter (P*_pck-2_pck-2::mCh*). The fusion protein was cytoplasmic, and it successfully rescued the spherical mitochondria in both *yq270* and *yq271* mutants to wild-type levels of mitochondrial tubules (Fig 1H). Collectively, these results suggest that loss of *pck-2* function causes mitochondrial abnormalities.

Humans have two PEPCK homologs, the cytosolic PCK1 (hPCK1) and mitochondrial PCK2 (hPCK2) (S1C, S1D and S2 Figs). We tested their rescue ability via transgenic expression of hPCK1 (P*_semo-1_hPCK1::gfp*) or hPCK2 with deletion of its N-terminal mitochondrion-targeting sequence (P*_semo-1_hPCK2(ΔMTS)::gfp*). Both proteins were distributed in the cytoplasm, and they mostly rescued the abnormal mitochondrial morphology in *C. elegans pck-2(ok2586)* mutants (Fig 1I). However, expression of the full-length hPCK2 (P*_semo-1_hPCK2::gfp*), which localized to mitochondria, failed to rescue the defective mitochondria in *C. elegans pck-2* mutants (S1E Fig). These results suggest that PCK activity is required in the cytoplasm to maintain mitochondrial morphology, and this function is evolutionarily conserved.

### *C. elegans* PCK-1 and PCK-2 act together to affect mitochondrial homeostasis

In addition to PCK2, *C. elegans* possesses an additional PEPCK, PCK-1 (S1B, S1C and S2 Figs). To investigate whether PCK-1 is also required for normal mitochondrial morphology, we analyzed *pck-1(ok2098)* deletion mutants (S1B Fig) and double mutants of *pck-1(ok2098)* with *pck-2(ok2586)*. *pck-1(ok2098)* animals exhibited spherical mitochondria in the hypodermis and body-wall muscles, and *pck-2;pck-1* double mutants displayed abnormal mitochondria, similar to single mutants of either *pck* gene (Fig 2A). Using transmission electron microscopy (TEM), we confirmed that mitochondria were spherical and the organization of cristae was similarly disrupted in both *pck-1(ok2098)* and *pck-2(ok2586)* single mutants as well as their double mutants (Fig 2B). Moreover, these *pck* single and double mutants exhibited similar reductions in oxygen consumption rate (OCR), ATP production and body bending (Fig 2C-2E), as well as shortened body lengths and lifespan (Fig 2F and 2G). These results suggest that *pck-1* function is required for normal mitochondrial morphology, and that *pck-1* acts in the same genetic pathway with *pck-2*. Using in vitro GST pull-down assays, we found that recombinant His_6_-PCK-2 directly interacted with GST-PCK-1 (Fig 2H). Transgenic PCK-1::GFP driven by the *pck-1* promoter (P*_pck-1_pck-1::gfp*), which localized to the cytoplasm, successfully rescued the spherical mitochondria in *pck-1(ok2098)* mutants but not *pck-2(ok2586)* mutants (Fig 2I). In comparison, transgenic PCK-2::GFP (P*_pck-2_pck-2::gfp*) fully rescued the spherical mitochondria in both *pck-1* and *pck-2* deletion mutants (Fig 2I). Taken together, these results suggest that PCK-1 and PCK-2 function together, and PCK-2 acts downstream of PCK-1 in maintaining normal mitochondrial structure and functions.

**Fig 2.**
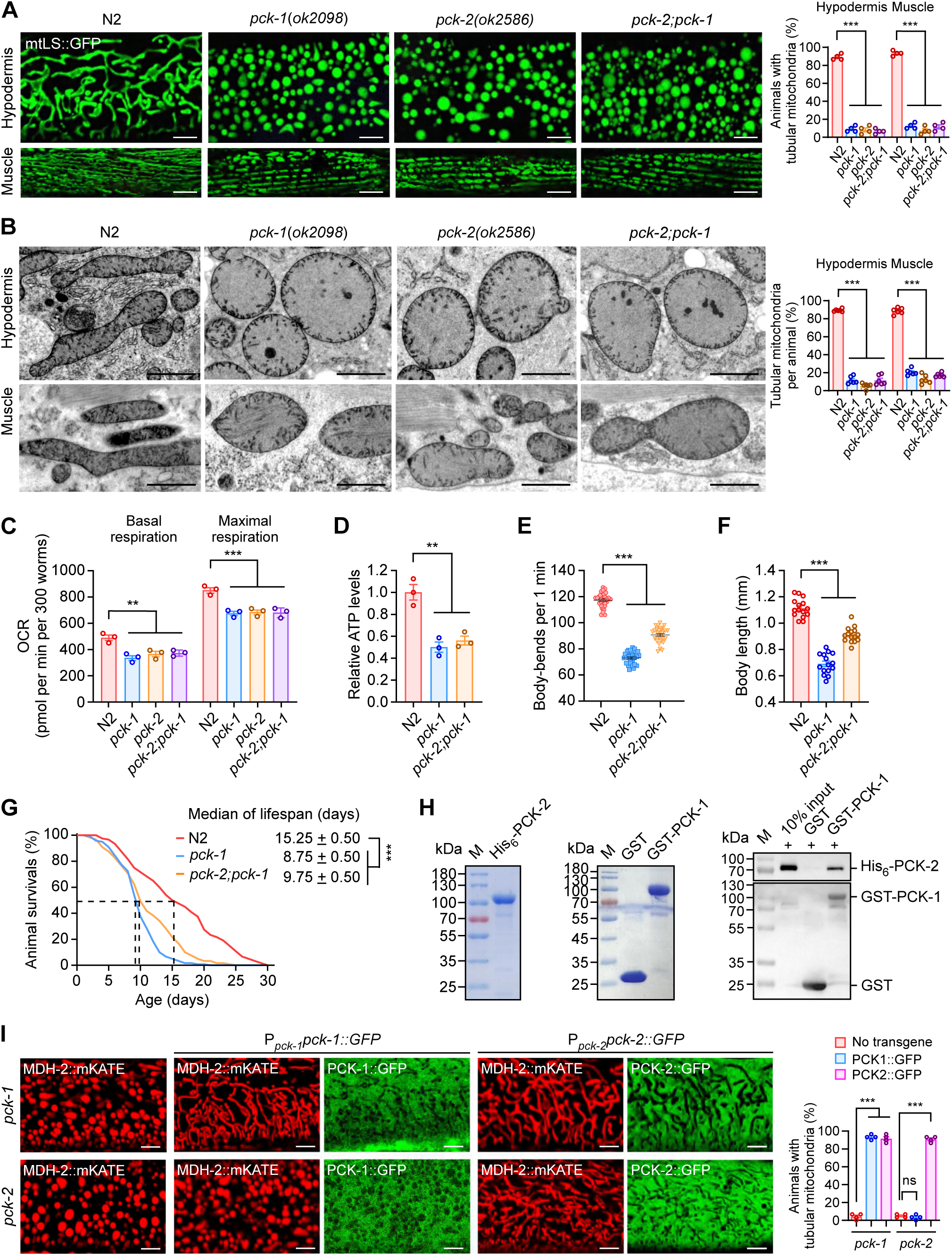
*C. elegans* PCK-1 and PCK-2 act together. **(A)** Representative images (left) and quantification (right) of mitochondria (labeled with MITO-GFP) in the hypodermis and muscle of N2, *pck-1(ok2098)* single mutants, *pck-2(ok2586)* single mutants and *pck-2;pck-1* double mutants at the adult stage (48 h post L4). Bars, 5 μm. A total of 120 animals were examined for each genotype in *n*= 4 independent experiments. **(B)** Representative TEM images (left) and quantification (right) of mitochondria in the hypodermis and muscle of N2, *pck-1*(*ok2098*), *pck-2*(*ok2586*), and *pck-2*(*ok2586*);*pck-1*(*ok2098*) animals at 48 h post L4 stage. Bars, 1 μm. A total of 600 mitochondria from *n*=6 animals were examined for each genotype. **(C-G)** OCR (300 animals for each genotype, *n*= 3 independent experiments) (C), relative ATP levels (300 animals for each genotype, *n*= 3 independent experiments) (D), body-bending rates (*n*= 30 animals for each genotype) (E), body length (n=15 animals for each genotype) (F) and lifespan of adult animals (*n*=120 animals for each genotype) (G). **(H)** PCK-1 and PCK-2 interact with one another. Left and middle: Coomassie blue staining of purified His_6_-PCK-2 and GST-PCK-1. Right: GST and GST-PCK-1 bound on glutathione sepharose beads were incubated with His_6_-PCK-2. After extensive washing, bound proteins were detected with His_6_ and GST antibodies. **(I)** Representative images (left) and quantification (right) of the rescuing effect on *pck-1(ok2098)* and *pck-2(ok2586)* mitochondria (labeled with MDH-2::mKATE) by transgenic PCK-1::GFP (P*_pck-1_pck-1::gfp*) or PCK-2::GFP (P*_pck- 2_pck-2::gfp*). A total of 120 animals were examined for each genotype in *n*= 4 independent experiments. Bars, 5 μm.For quantifications, data points represent (mean ± s.e.m.). *P* values were determined using one-way ANOVA in (A**-**G and I). \*\*\**P*<0.001; \*\**P* < 0.01; \**P* < 0.05; ns, not significant (*P* > 0.05).

### Loss of *pck-1* and/or *pck-2* leads to OAA buildup that impairs mitochondria

As PEPCKs, PCK-1 and PCK-2 act in the cytoplasm to convert OAA to PEP (Fig 3A and 3B). We thus analyzed OAA levels in single mutants and double mutants of *pck-1* and *pck-2*. Indeed, OAA levels were significantly increased to similar levels in these single and double mutants (Fig 3C). Further analysis indicated that OAA was increased in both cytosol and mitochondria (Fig 3D). Thus, loss of *pck-1* and/or *pck-2* probably blocks the conversion of OAA to PEP, leading to OAA accumulation in both cytosol and mitochondria. To determine whether OAA accumulation accounts for the abnormal mitochondria in *pck* mutants, we generated double mutants of *pck-1* or *pck-2* with *got-1.2(yq163)*, a mutation causing a premature stop at W312 in the GOT-1.2 transaminase that converts Asp to OAA (S3A and S3B Fig). Unlike the spherical mitochondria in *pck* mutants, single mutants of *got-1.2(yq163)* and double mutants with *pck-1* or *pck-2* displayed very well-connected tubular mitochondria, which were confirmed by TEM analysis (Fig 3E and 3F). Importantly, these mutants had significantly lower levels of cytosolic and mitochondrial OAA than *pck-1* and *pck-2* mutants (Fig 3G), and their ATP levels were significantly higher than in *pck-1* and *pck-2* mutants (Fig 3H). Collectively, these findings suggest that the aberrantly accumulated OAA is responsible for mitochondrial abnormalities in *pck* mutants.

**Fig 3.**
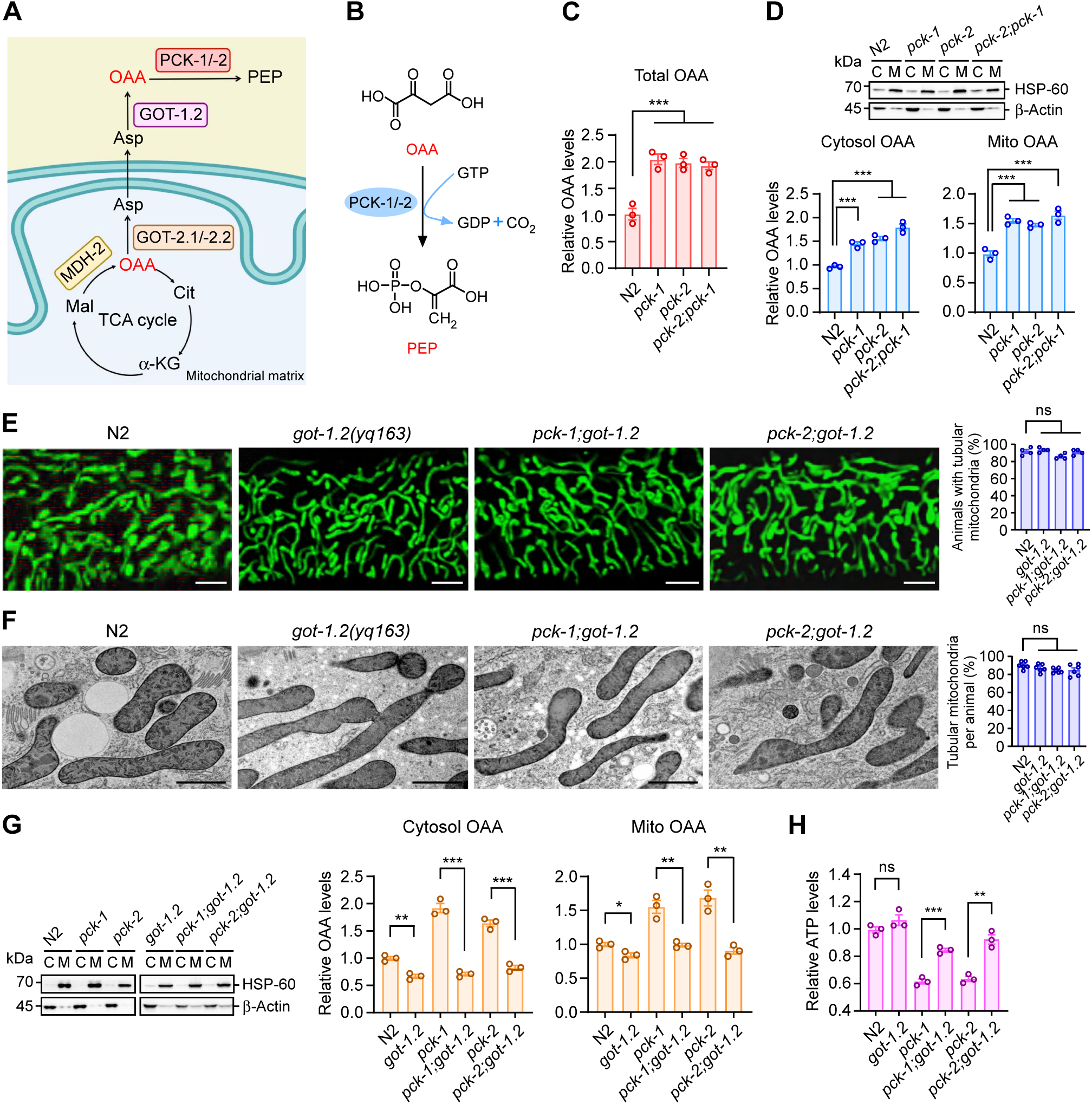
The increase in OAA levels causes the abnormal mitochondrial phenotype in *pck-1* and *pck-2* mutants. **(A)** Graphical summary of OAA metabolism in the cytosol and mitochondria in *C. elegans*. α-KG, α-ketoglutarate; Asp, aspartate; Cit, citrate; Mal, malate; MDH-2, malate dehydrogenase-2; OAA, oxaloacetate. **(B)** Schematic showing the reaction of OAA conversion to PEP by PCK-1/-2. **(C** and **D)** Relative OAA levels in total worm lysates (C) and cytosolic and mitochondrial fractions (D) from animals with the indicated genotypes. In the top panel in (D), western blotting of subcellular fractions (C: cytosol; M: mitochondria) is shown with antibodies against HSP-60 and β-Actin to indicate mitochondrial fractions and cytosolic fractions, respectively. In the bottom panels, relative cytosolic and mitochondrial OAA levels are shown on the left and right, respectively. *n*= 3 independent experiments. **(E)** Representative images (left) and quantification (right) of mitochondria (labeled with MITO-GFP) in the hypodermis of N2, *got-1.2*(*yq163*), *pck-1*(*ok2098*); *got-1.2*(*yq163*) and *pck-2*(*ok2586*)*;got-1.2*(*yq163*) animals at 48 h post L4. A total of 120 animals were examined for each genotype in *n*= 4 independent experiments. Bars, 5 μm. **(F)** Representative TEM images (left) and quantification (right) of mitochondria in the hypodermal cells in adult N2, *got-1.2*(*yq163*), *pck-1*(*ok2098*)*;got-1.2*(*yq163*) and *pck-2*(*ok2586*)*;got-1.2*(*yq163*) animals. Bars, 1 μm. A total of 600 mitochondria from *n*=6 animals were examined for each genotype. **(G)** Relative OAA levels in cytosolic and mitochondrial fractions from animals with the indicated genotypes. Western blotting of subcellular fractions (C: cytosol; M: mitochondria) is shown on the left with antibodies detecting HSP-60 and β-Actin to indicate mitochondrial fractions and cytosolic fractions, respectively. Relative cytosolic and mitochondrial OAA levels are shown in the middle and on the right, respectively. *n*= 3 independent experiments. **(H)** Relative ATP levels in animals with the indicated genotypes. 300 animals were analyzed for each genotype, *n*= 3 independent experiments. For quantifications, data points represent (mean ± s.e.m.). *P* values were determined using one-way ANOVA in (C-F) and the two-tailed unpaired Student’s *t*-test in (G and H). \*\*\**P*<0.001; \*\**P* < 0.01; \**P* < 0.05; ns, not significant (*P* > 0.05).

### Mitochondrial accumulation of OAA causes mitochondrial impairment

While PCK-2 and PCK-1 localize and function in the cytoplasm (Fig 2I), their deficiencies lead to OAA elevation in mitochondria (Fig 3C and 3D). To determine whether the elevated mitochondrial OAA directly causes mitochondrial abnormalities, we investigated mutants of *got-2.1* and *got-2.2*, which encode mitochondrial glutamic-oxaloacetic transaminases that convert OAA to Asp in mitochondria[15] (S3C-S3F Fig). As expected, *got-2.1(gk109644)* and *got-2.2(yq341)* mutants had significantly higher mitochondrial OAA levels than the wild type (Fig 4A). Consistent with the elevated mitochondrial OAA, *got-2.1(gk109644)* and *got-2.2(yq341)* animals exhibited spherical mitochondria with disrupted cristae, as in *pck-1* and *pck-2* mutants (Fig 4B and 4C). In addition, mitochondrial functions were impaired in these mutant animals, manifesting as significantly reduced ATP levels, body-bending rates, OCRs and lifespans (Fig 4D-4G).

**Fig 4.**
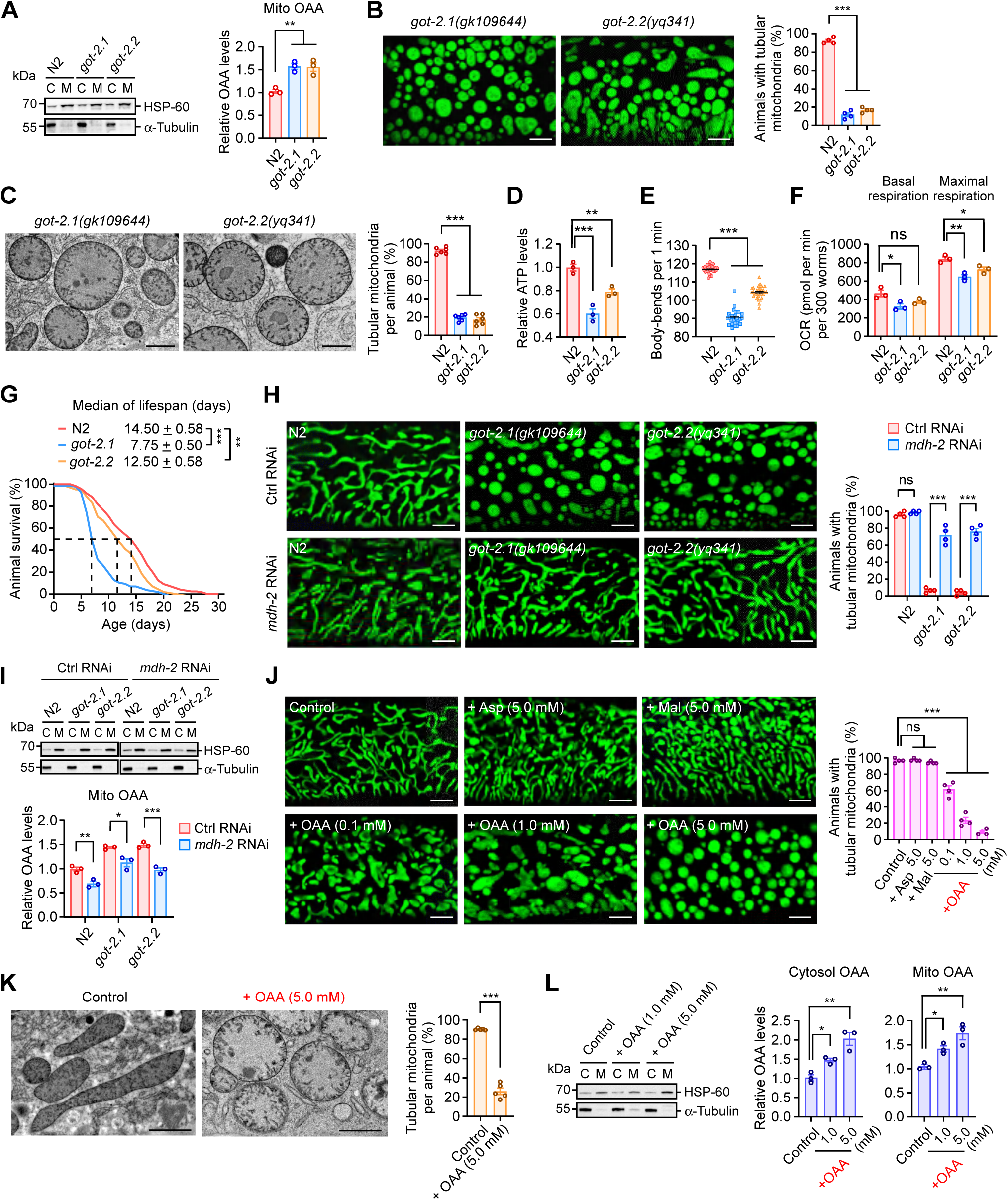
Mitochondrial OAA accumulation impairs mitochondrial structure. **(A)** Relative mitochondrial OAA levels in N2, *got-2.1(ok109644)* and g*ot-2.2(yq341)* animals with the indicated genotypes. Western blotting of subcellular fractions (C: cytosol; M: mitochondria) is shown on the left with antibodies detecting HSP-60 and α-Tubulin to indicate mitochondrial fractions and cytosolic fractions, respectively. *n*= 3 independent experiments. **(B)** Representative images (left) and quantification (right) of mitochondria (labeled with MITO-GFP) in the hypodermis of *got-2.1*(*gk109644*) and *got-2.2*(*yq341*) animals. Bars, 5 μm. A total of 120 animals were examined for each genotype in *n*= 4 independent experiments. **(C)** Representative TEM images (left) and quantification (right) of mitochondria in hypodermal cells in adult *got-2.1*(*gk109644*) and *got-2.2*(*yq341*) animals (48 h post L4). Bars, 1 μm. A total of 600 mitochondria from *n*=6 animals were examined for each genotype. **(D-G)** Relative ATP levels (300 animals for each genotype, *n*= 4 independent experiments) (D), body-bending rates (*n*= 30 animals for each genotype) (E), OCR (300 animals for each genotype, *n*= 3 independent experiments) (F), survival curves of adult animals (*n*= 120 animals for each genotype) (G) in animals with the indicated genotype. **(H)** Representative images (left) and quantification (right) of mitochondria (labeled with MITO-GFP) in the hypodermis of the indicated animals treated with control (Ctrl) RNAi or *mdh-2* RNAi. RNAi was performed using RNAi-competent OP50 *E. coli*. A total of 120 animals were examined for each genotype in *n*= 4 independent experiments. Bars, 5 μm. **(I)** Relative OAA levels in mitochondrial fractions of animals with the indicated genotypes. Western blotting of subcellular fractions (C: cytosol; M: mitochondria) is shown on the left with antibodies detecting HSP-60 and α-Tubulin to indicate mitochondrial fractions and cytosolic fractions, respectively. *n*= 3 independent experiments. **(J)** Representative images (left) and quantification (right) of mitochondria (labeled with MITO-GFP) in the hypodermis in adult N2 animals treated Asp, Mal, and OAA at the indicated concentrations. A total of 120 animals were examined for each treatment in *n*= 4 independent experiments. Bars, 5 μm. **(K)** TEM images (left) and quantification (right) of mitochondria in the hypodermis of adult N2 animals treated without and with OAA (5.0 mM). A total of 600 mitochondria from 5 animals were examined for each group. Bars, 1 μm. **(L)** Relative OAA levels in cytosolic and mitochondrial fractions of N2 animals treated without and with OAA. Western blotting of subcellular fractions (C: cytosol; M: mitochondria) is shown on the left with antibodies against HSP-60 and α-Tubulin to indicate mitochondrial fractions and cytosolic fractions, respectively. Relative cytosolic and mitochondrial OAA levels are shown in the middle and on the right, respectively. *n*= 3 independent experiments. For quantifications, data points represent (mean ± s.e.m.). *P* values were determined using one-way ANOVA in (A-G, J and L) or the two-tailed unpaired Student’s *t*-test in (H, I and K). \*\*\**P*<0.001; \*\**P* < 0.01; \**P* < 0.05; ns, not significant (*P* > 0.05).

Mitochondrial OAA is a pivotal metabolic intermediate primarily produced by malate dehydrogenase 2 (MDH-2) in the TCA cycle (Fig 3A). We sought to reduce OAA levels with RNAi knockdown of *mdh-2*, the gene that encodes mitochondrial MDH. *mdh-2* RNAi significantly reduced mitochondrial OAA levels in *got-2.1(gk109644)* animals and *got-2.2(yq341)* animals (Fig 4I). Consistent with this, *mdh-2* RNAi-treated *got-2.1(gk109644)* mutants and *got-2.2(yq341)* mutants displayed well-connected tubular mitochondria (Fig 4H). These findings suggest that aberrantly accumulated OAA in mitochondria causes mitochondrial impairment. To corroborate this conclusion, we fed N2 animals OAA and found that it caused mitochondrial fragmentation in a concentration-dependent manner (Fig 4J). In contrast, similar supplementation of Asp or Mal did not induce an obvious change in mitochondrial morphology (Fig 4J). By TEM analysis, we found that OAA treatment led to disruption of mitochondrial cristae, as observed in *pck-1/-2* and *got-2.1/-2.2* mutants (Fig 4K). Importantly, both cytosolic and mitochondrial OAA levels were significantly elevated following exogenous OAA supplementation (Fig 4L). Taken together, these results suggest that excessive accumulation of OAA in mitochondria causes mitochondrial impairment and dysfunction.

### Reinforced expression of IMMT-1 or CHCH-3 ameliorates mitochondrial defects in *C. elegans* mutants with OAA accumulation

We hypothesized that aberrantly accumulated OAA in mitochondria probably binds to and interferes with proteins that are required for cristae organization and mitochondrial functions. These include EAT-3/Opa1, FZO-1/Mfn1/2, the MICOS complex, respiratory chain complexes, and the ATP synthase complex[3, 4, 36]. To investigate this possibility, we reinforced the expression of individual proteins in *pck-1/-2* mutants (S4 Fig). Our results revealed that reinforced expression of IMMT-1/MIC60, which is capable of membrane binding and shaping[37, 38], and CHCH-3/MIC19, which promotes IMMT-1/MIC60 activities[39, 40], mostly restored the spherical mitochondria to tubular morphology in *pck-1/-2* single and double mutants, and in *got-2.1* and *got-2.2* mutants (Figs 5A and S5A). TEM analysis further confirmed that the organization of cristae was restored to normal (Figs 5B and S5B). Overexpression of other components involved in cristae organization did not have similar effects (S4 Fig). In addition, while exogenously supplied OAA induced mitochondrial fragmentation in the wild type, it did not cause a marked change of mitochondrial morphology in animals overexpressing either IMMT-1 or CHCH-3 (Fig 5C). Thus, reinforced expression of IMMT-1 and CHCH-3 antagonizes OAA-induced mitochondrial abnormalities. These results collectively suggest that OAA probably impairs mitochondria through IMMT-1 and/or CHCH-3.

**Fig 5.**
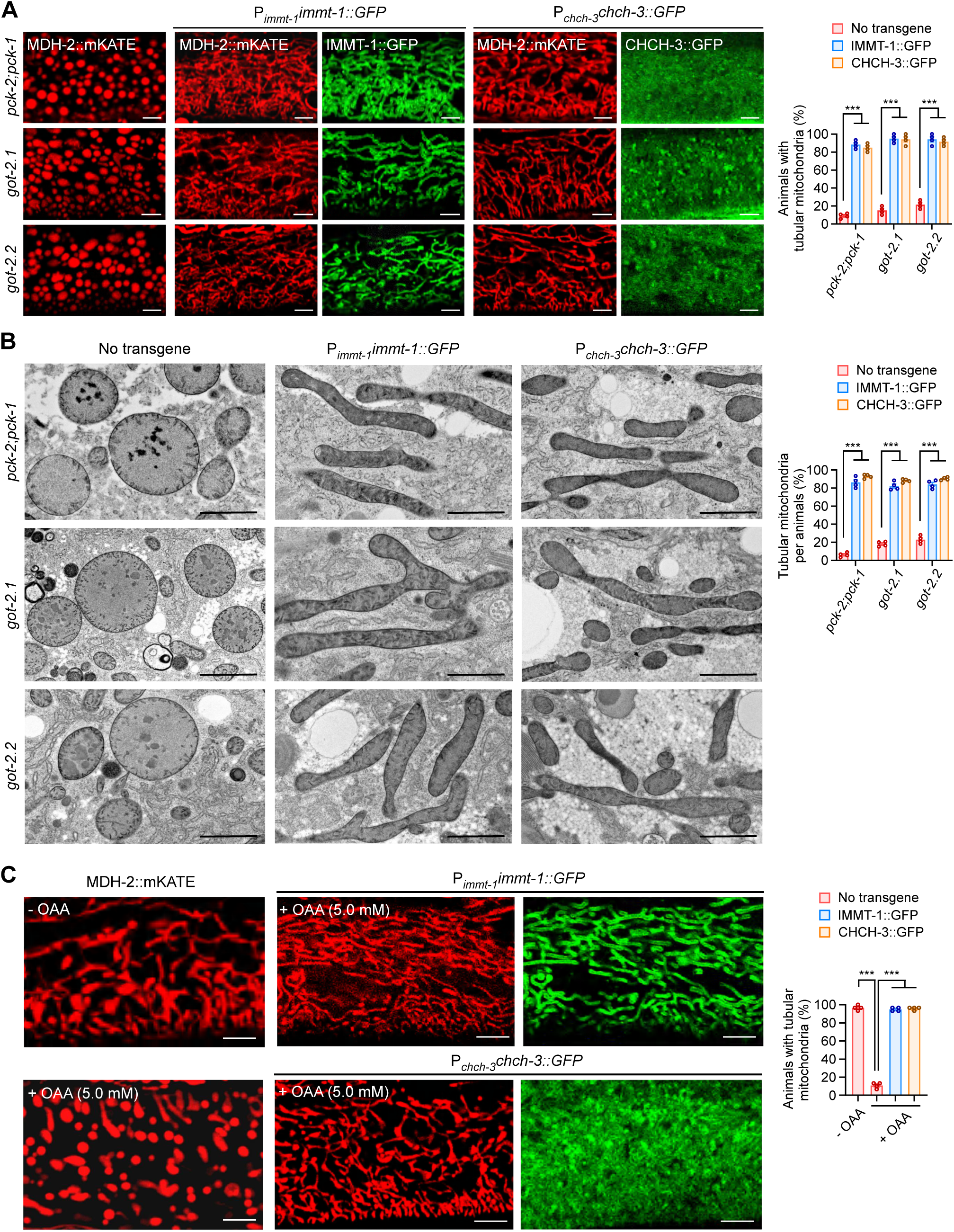
Reinforced expression of IMMT-1 and CHCH-3 restores tubular mitochondrial morphology in OAA-accumulating mutants. **(A)** Representative images (left) and quantification (right) of mitochondria (labeled with MDH-2::mKATE) in the hypodermis of *pck-2*(*ok2586*)*;pck-1*(*ok2098*), *got-2.1*(*gk109644*) and *got-2.2*(*yq341*) mutants without or with overexpression of IMMT-1::GFP (P*_immt-1_immt-1::gfp*) or CHCH-3::GFP (P*_chch-3_chch-3::gfp*). A total of 120 animals were examined for each genotype in *n*= 4 independent experiments. Bars, 5 μm. **(B)** Representative TEM images (left) and quantification (right) of mitochondria in the hypodermis of *pck-2*(*ok2586*)*;pck-1*(*ok2098*), *got-2.1*(*gk109644*) and *got-2.2*(*yq341*) mutants without or with overexpression of IMMT-1::GFP (P*_immt-1_immt-1::gfp*) or CHCH-3::GFP (P*_chch-3_chch-3::gfp*). A total of 400 mitochondria from *n*= 4 animals were examined for each genotype. Bars, 1 μm. **(C)** Representative images (left) and quantification (right) of mitochondria (labeled with MDH-2::mKATE) in OAA (5.0 mM)-treated N2 animals without or with overexpression of IMMT-1::GFP (P*_immt-1_immt-1::gfp*) or CHCH-3::GFP (P*_chch-3_chch-3::gfp*). A total of 120 animals were examined for each genotype in *n*= 4 independent experiments. Bars, 5 μm. For quantifications, data points represent (mean ± s.e.m.). *P* values were determined using one-way ANOVA in (A and B) or two-way ANOVA in (C). \*\*\**P*<0.001; \*\**P* < 0.01; \**P* < 0.05; ns, not significant (*P* > 0.05).

### OAA binds to CHCH-3 and inhibits its function of promoting IMMT-1-dependent membrane shaping

To determine if OAA acts through IMMT-1 and/or CHCH-3, we first examined their expression levels in *pck* mutants that accumulate OAA. Neither mRNA nor protein levels of IMMT-1, CHCH-3, or several other MICOS components (F54A3.5(MIC10), MOMA-1(MIC27), W04C9.2(MIC13)) were changed in single or double *pck* mutants (S6A and S6B Fig). This suggests that OAA does not affect mitochondria by altering the expression of these genes. We then examined whether OAA binds with IMMT-1 and CHCH-3. In microscale thermoelectrophoresis (MST) assays, OAA bound well to purified His_6_-CHCH-3 (*K*d=2.69 ± 0.57 μM); however, no obvious OAA binding with His_6_-IMMT-1 was detected (Fig 6A). Further analysis with the AutoDock Vina 1.2.0 algorithm[41, 42] suggests that OAA probably binds to CHCH-3 at the C-terminal CHCH domain by forming hydrogen bonds with Asn129, Ala153, and Glu160 residues (Fig 6B). Consistent with this prediction, CHCH-3(N129A) and CHCH-3(E160A) mutants did not bind with OAA in MST assays (Fig 6C). Thus, the interaction of OAA with CHCH-3 is specific.

**Fig 6.**
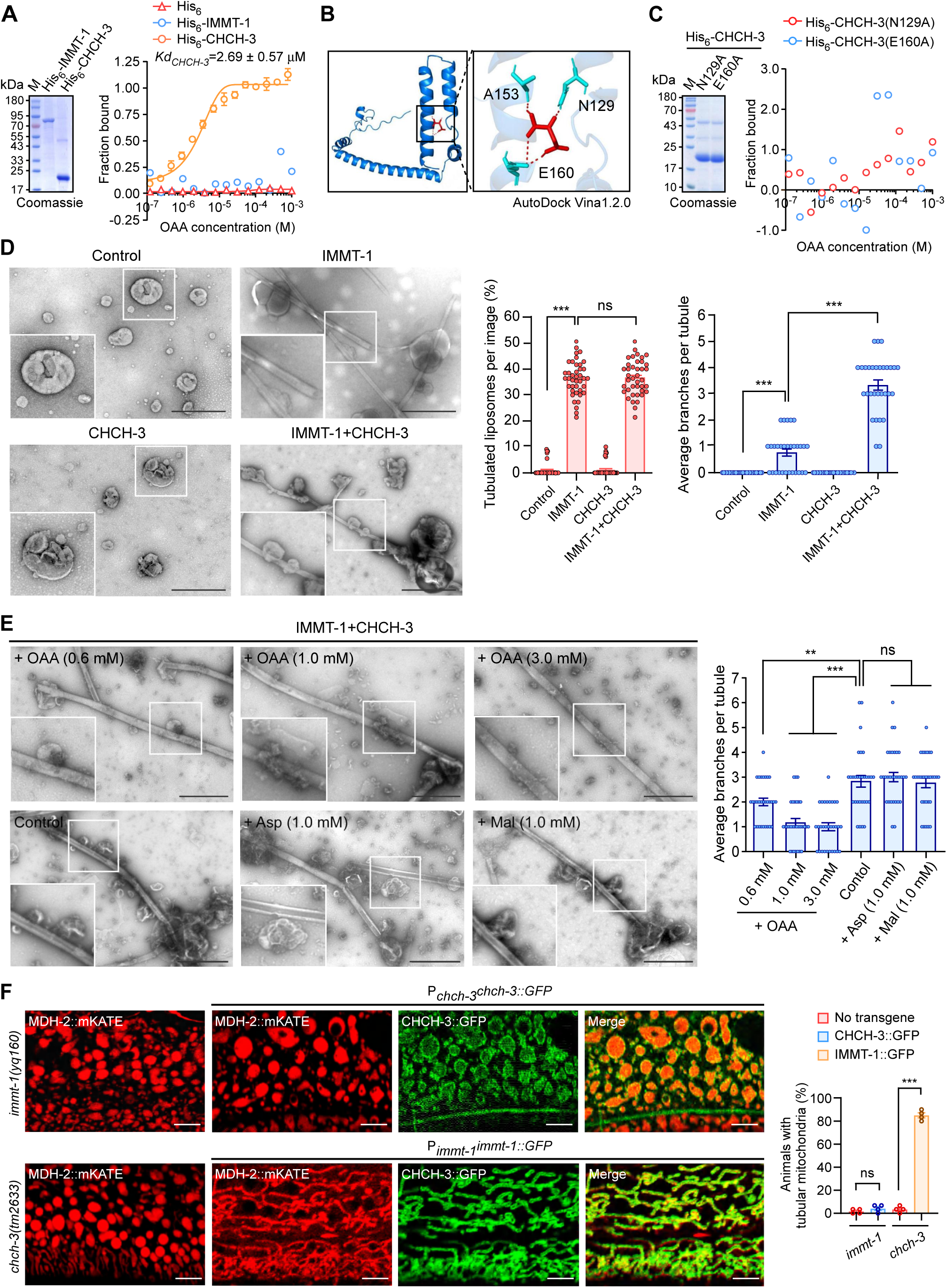
OAA binds to CHCH-3 and inhibits IMMT-1-dependent membrane remodeling. **(A)** Binding curves, determined by microscale thermoelectrophoresis (MST), of OAA with His_6_-IMMT-1 and His_6_-CHCH3. Coomassie blue staining of purified His_6_-IMMT-1 and His_6_-CHCH-3 is shown on the left. *n*= 3 independent experiments. **(B)** Prediction of OAA binding sites in CHCH-3 with the AutoDock Vina 1.2.0 software. The framed region in the left image is magnified in the right image to show the hydrogen bonds (indicated with red dashed lines) formed between OAA (red) and N129, A153 and E160 residues (blue) in CHCH3. **(C)** MST binding curves of OAA with His_6_-CHCH-3(N129A) or His_6_-CHCH-3(E160A) point mutants. Coomassie blue staining of purified His_6_-CHCH-3 with the indicated point mutations is shown on the left. **(D)** Representative TEM images (left) and quantifications (right) of membrane shaping of liposomes by IMMT-1, CHCH-3, and CHCH-3 with IMMT-1. Bars, 1 μm. ≥30 liposomes were scored in each group. **(E)** Representative TEM images (left) and quantification (right) of membrane shaping of liposomes by CHCH-3 together with IMMT-1 in the presence of OAA (0.6, 1.0 and 3.0 mM), Asp (1.0 mM), and Mal (1.0 mM). Bars, 1 μm. ≥30 liposomes were scored in each treatment. **(F)** Representative images (left) and quantification (right) of the rescue of *immt-1(yq160*) mitochondria by transgenic CHCH-3::GFP (P*_chch-3_chch-3::gfp*) or the rescue of *chch-3*(*tm2633*) mitochondria by transgenic IMMT-1::GFP (P*_immt-1_immt-1::gfp*). Mitochondria were labeled with MDH-2::mKATE. Bars, 5 μm. A total of 120 animals were examined in *n*= 4 independent groups for each genotype. For quantifications, data points represent (mean ± s.e.m.). *P* values were determined using one-way ANOVA in (E) or the two-tailed unpaired Student’s *t*-test in (D and F). ***, *P*<0.001; \*\**P* < 0.01; \**P* < 0.05; ns, not significant (*P* > 0.05).

In the fungus *Chaetomium thermophilum*, Mic19, the homolog of *C. elegans* CHCH-3, interacts with Mic60 and promotes Mic60-dependent membrane shaping[39, 40]. To test this interaction in *C. elegans*, we knocked in GFP at the C-terminus of IMMT-1, then performed co-immunoprecipitation assays. We found that endogenous CHCH-3 was indeed precipitated with endogenous-level IMMT-1::GFP from cell lysates of both wild type (N2) and *pck* mutants (S6C Fig). In in vitro GST pull-down assays, increasing the OAA concentration did not markedly affect the interaction of CHCH-3 with IMMT-1 (S6D Fig). Together these results suggest that the aberrantly accumulated OAA in *pck-1* and *pck-2* mutants probably does not affect the interaction of CHCH-3 with IMMT-1. We next investigated whether OAA interferes with IMMT-1-dependent membrane shaping using liposome assays. We prepared liposomes and incubated them with His_6_-IMMT-1 or His_6_-CHCH-3. His_6_-IMMT-1 induced obvious tubulation of liposomes, while His_6_-CHCH-3 did not have the same effect (Fig 6D). Notably, when both His_6_-CHCH-3 and His_6_-IMMT-1 were together incubated with liposomes, they generated membrane tubules with branches or bouton-like structures (Fig 6D). These results suggest that CHCH-3 acts through IMMT-1 to promote membrane shaping, which is consistent with previous reports that fungal Mic19 promotes Mic60-dependent membrane shaping[39, 40]. In the presence of OAA, the branching of membrane tubules by the coordinated action of His_6_-CHCH-3 with His_6_-IMMT-1 was inhibited in a concentration-dependent manner, while Asp and Mal did not have marked inhibitory effects (Fig 6E). These results suggest that OAA binds to CHCH-3 and prevents it from promoting IMMT-1-dependent membrane shaping. Taking these results together with the findings that reinforced expression of either CHCH-3 or IMMT-1 ameliorated the mitochondrial defects in animals accumulating OAA (Figs 5A and S5A), we reasoned that *immt-1* probably acts downstream of *chch-3* in the maintenance of mitochondrial homeostasis. To test this possibility, we overexpressed IMMT-1::GFP in *chch-3(tm26333)* mutants, and CHCH-3::GFP in *immt-1(yq160)* mutants. Overexpressing IMMT-1::GFP mostly restored the mitochondria in *chch-3* mutants to wild-type tubular morphology, while overexpressing *chch-3* failed to ameliorate the mitochondrial fragmentation in *immt-1* mutants (Fig 6F). Altogether, these findings provide genetic evidence that CHCH-3 acts through IMMT-1, and suggest a mechanism in which excessively accumulated OAA disrupts mitochondrial homeostasis by inhibiting CHCH-3 function.

### OAA accumulation impairs mitochondria in mammalian cells

We finally investigated whether OAA accumulation similarly causes mitochondrial impairment in mammalian cells. 3-MPA (3-mercaptopicolinic acid) is an inhibitor of human PCK1 and PCK2 [43, 44]. In U2OS and HeLa cells, 3-MPA treatment induced obvious mitochondrial fragmentation in a concentration-dependent manner (Figs 7A and S7). Consistent with this, 3-MPA led to a significant elevation of total and mitochondrial OAA levels in U2OS cells (Fig 7B). These results suggest that inhibition of hPCK1 and hPCK2 causes mitochondrial accumulation of OAA, which induces mitochondrial fragmentation as in *C. elegans*. To consolidate this point, we treated cells with OAA. As in *C. elegans*, exogenously supplied OAA led to mitochondrial fragmentation in U2OS and HeLa cells (Figs 7C and S7), and the total cellular and mitochondrial OAA levels were significantly increased (Fig 7D). TEM analysis further revealed that both 3-MPA and OAA disrupted the structures of mitochondrial cristae to similar levels (Fig 7E). In contrast, adding Asp and Mal to U2OS and HeLa cells did not induce mitochondrial fragmentation (Figs 7C and S7). To investigate whether OAA acts through MIC19 and MIC60 to impair cristae in mammalian cells, we overexpressed MIC19-EGFP or MIC60-EGFP in U2OS cells and treated them with 3-MPA or OAA. Cells overexpressing MIC19-EGFP or MIC60-EGFP were resistant to mitochondrial fragmentation induced by 3-MPA or OAA (Fig 7F and 7G). Taken together, these results suggest that OAA accumulation damages mitochondria in mammalian cells in a similar way to that in *C. elegans*.

**Fig 7.**
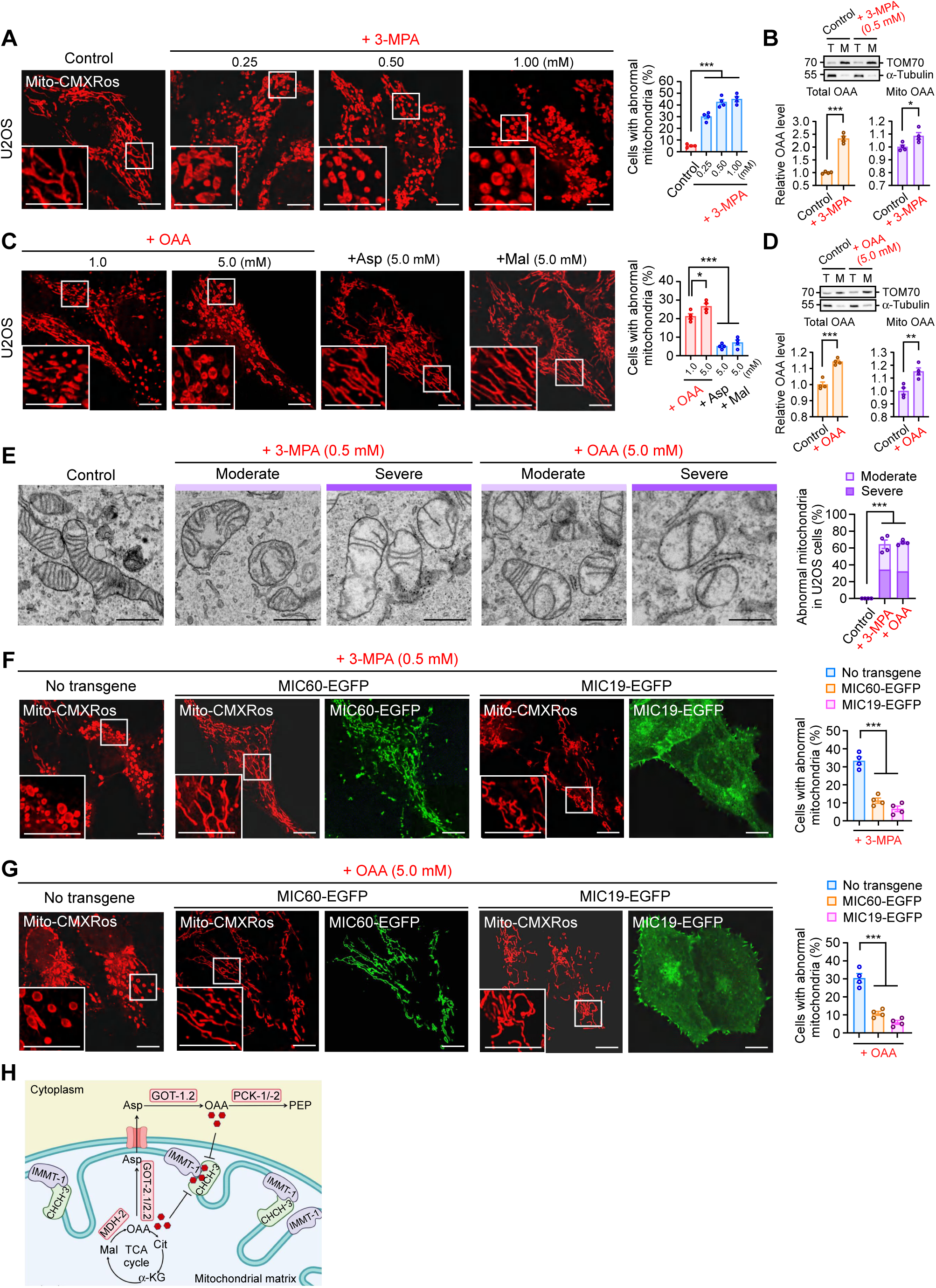
OAA induces mitochondrial defects through MIC60 and MIC19 in mammalian cells. **(A)** Representative images (left) and quantification (right) of abnormal mitochondria in U2OS cells treated with 3-MPA at the indicated concentrations. Mitochondria were labeled with Mito-CMXRos. Boxed regions are magnified (2.5×) in the bottom left. Bars, 10 μm. A total of ≥240 cells in n= 4 independent repeats were scored in each group. **(B)** Relative levels of total cellular and mitochondrial OAA in U2OS cells treated with or without 3-MPA (0.5 mM). Western blotting of subcellular fractions (T: total; M: mitochondria) is shown at the top with antibodies against TOM70 and α-Tubulin to indicate mitochondrial fractions and cytosolic fractions, respectively. *n*= 4 independent repeats. **(C)** Images (left) and quantification (right) of mitochondria in U2OS cells treated with OAA (1.0 and 5.0 mM), Asp (5.0 mM), or Mal (5.0 mM). Mitochondria were stained with Mito-CMXRos. Boxed regions are magnified (2.5×) in the bottom left. A total of ≥240 cells were scored in *n*= 4 repeats for each treatment. **(D)** Relative levels of total cellular and mitochondrial OAA in U2OS cells treated with or without OAA (5.0 mM). Western blotting of subcellular fractions (T: total; M: mitochondria) is shown at the top with antibodies against TOM70 and α-Tubulin to indicate mitochondrial fractions and cytosolic fractions, respectively. *n*= 4 independent repeats. **(E)** TEM images (left) and quantification (right) of abnormal mitochondria in U2OS cells treated with 3-MPA (0.5 mM) or OAA (5.0 mM). Mitochondria were categorized into “moderate” and “severe” impairment groups. A total of ≥ 400 mitochondria were analyzed in *n*= 4 cells for each treatment. Bars, 1 μm. **(F** and **G)** Representative images (left) and quantification (right) of mitochondria in U2OS cells treated with 3-MPA (F) or OAA (G) without or with overexpression of MIC19-EGFP or MIC60-EGFP. Mitochondria were stained with Mito-CMXRos. Boxed regions are magnified (2.5×) in the bottom left. Bars, 10 μm. A total of ≥ 240 cells were analyzed in *n*=4 repeats. **(H)** Graphic summary of OAA-induced mitochondrial damage. Loss-of-function mutations in cytosolic PCK-1/-2 or mitochondrial GOT-2.1/-2.2 cause buildup of OAA in mitochondria. OAA binds to CHCH-3 and inhibits its function of promoting IMMT-1-dependent membrane remodeling, leading to impaired organization of mitochondrial cristae. For all quantifications, data points represent (mean ± s.e.m.). *P* values were determined using one-way ANOVA in (A, C, E, F and G) or the two-tailed unpaired Student’s *t*-test in (B and D). \*\*\**P* < 0.001; \*\**P* < 0.01; \**P* < 0.05; ns, not significant (*P* > 0.05).

## Discussion

In this study, we demonstrated that aberrant accumulation of OAA disrupts mitochondrial architecture and functions. Using unbiased genetic screening, we revealed that mutations of *C. elegans pck-2* cause mitochondrial fragmentation and impairment of mitochondrial functions. We further found that *C. elegans* PCK-1 and PCK-2 act together in OAA metabolism, and their loss leads to buildup of both total and mitochondrial OAA, which severely damage mitochondrial architecture and functions. Genetic depletion of the cytosolic GOT-1.2, which catalyzes the conversion of Asp to OAA, suppresses the elevation of OAA levels and consequently mitochondrial impairment, suggesting that OAA is responsible for mitochondrial damage in *pck-1/-2* mutants. Similar to *pck-1/-2* mutations, mutations of mitochondrial GOT-2.1/-2.2, which catalyze the conversion of OAA to Asp in mitochondria, cause accumulation of mitochondrial OAA, mitochondrial fragmentation and disorganized mitochondrial architecture. Genetic inhibition of mitochondrial MDH-2, the enzyme that converts Mal to OAA, suppresses the OAA accumulation and mitochondrial impairment in *got-2.1/-2.2* mutants. This provides further evidence that OAA accumulation in mitochondria directly disrupts mitochondrial structure and functions. Moreover, exogenously supplied OAA induces significant elevation of total and mitochondrial OAA levels, leading to mitochondrial damage as seen in *C. elegans* mutants of *pck-1/-2* and *got-2.2/-2.2.* In mammalian cells, chemical inhibition of PCK1/2 or direct supply of OAA similarly induces mitochondrial impairment. Altogether, these findings uncover the adverse effects of excessive OAA accumulation on mitochondrial structure and functions (Fig 7H).

Our results demonstrated that OAA binds to CHCH-3/MIC19 to inhibit its function of promoting IMMT-1/MIC60-dependent membrane shaping. In support of this conclusion, reinforced expression of either CHCH-3/MIC19 or IMMT-1/MIC60 markedly ameliorated the mitochondrial defects in OAA-accumulating *C. elegans* animals and mammalian cells. The MICOS complex, composed of the MIC60 and MIC10 subcomplex, is essential for formation of the architecture of cristae[3, 38, 45, 46]. In the MIC60 subcomplex, MIC60/IMMT-1 binds and shapes membranes, while other components, including MIC19/CHCH-3, probably promote the membrane shaping activity of MIC60[39, 40]. Whereas reinforced expression of either *immt-1*/MIC60 or *chch-3*/MIC-19 rescued the defective mitochondrial structure in OAA-accumulating mutants, our in vitro analysis revealed that OAA only binds to CHCH-3 and not IMMT-1. Further analysis indicated that OAA binds to the CHCH domain of CHCH-3, which is required to promote IMMT-1 activities[40]. Thus, the OAA binding likely perturbs the function of the CHCH domain, though further structural investigation is needed to demonstrate the in-depth mechanism. These results, together with the findings that reinforced IMMT-1 expression bypassed the mitochondrial defects in *chch-3* mutants, but not vice versa, provide both in vitro and in vivo evidence that CHCH-3 acts through IMMT-1 to regulate the formation and maintenance of mitochondrial cristae. The non-covalent binding of OAA to CHCH-3 in this study, together with our previous findings that 3-hydoxypropionate (3-HP) binds to and inhibits IMMT-1/MIC60[10], suggests that the MICOS complex, and probably other factors required for maintenance of cristae organization, are susceptible to “attack” from excessively accumulated metabolic intermediates. It will be important to investigate how non-covalent inhibition and covalent modifications of cristae regulators by metabolic intermediates contribute differentially to mitochondrial damage or dysfunction in diverse organic acidemias/acidurias.

OAA is a key intermediate in the TCA cycle and multiple biochemical reactions, and therefore its concentration and availability determine the flow of the TCA cycle and the metabolic homeostasis of the cell. OAA was found to inhibit succinate dehydrogenase (SDH) activity[47, 48], competitively inhibit lactate dehydrogenase A (LDHA)[12] and promote antiviral innate immune responses[49]. Our findings that OAA inhibits the MICOS complex to damage mitochondria thus uncover a previously unrecognized effect of aberrantly accumulated OAA. It remains to be investigated whether OAA accumulation also disturbs the function of additional mitochondrial proteins. Disruption of OAA metabolism is found in multiple human diseases, including PEPCK deficiency and DEE82 (developmental and epileptic encephalopathy 82). Mutations of human cytosolic PCK1, the homolog of *C. elegans* PCK-2, lead to PEPCK deficiency, a disorder manifesting as neonatal hypoglycemia, acute liver failure, lactic acidosis and hyperammonemia. PEPCK deficiency patients also show progressive multisystem damage, hypotonia, developmental delay with seizures, spasticity, lethargy, microcephaly and cardiomyopathy, and they eventually fail to thrive[32–34]. While this disease is generally considered as resulting from defective gluconeogenesis, our findings suggest it is necessary to take into account that the pathological mechanism includes mitochondrial damage caused by excessive OAA accumulation. Similarly, mutations of human mitochondrial GOT2, the homolog of C. *elegans* GOT-2.1/-2.2 that catalyze the conversion of OAA to Asp, result in DEE82, an autosomal recessive mitochondriopathy manifesting as early-onset metabolic epileptic encephalopathy, characterized by hypotonia, feeding difficulties, and global developmental delay[15, 50, 51]. Our findings in this study thus provide important mechanistic insights into the mitochondrial damage in DEE82. More importantly, our genetic and metabolic analyses reveal that inhibiting cytosolic GOT1 and mitochondrial MDH2 could reduce the accumulation of OAA and alleviate mitochondrial damage, providing potential therapeutic targets for ameliorating these diseases.

## Materials and methods

### *C. elegans* strains and genetics

*C. elegans* cultures and genetic analysis were performed with standard procedures. The Bristol N2 strain was used as the wild type. *pck-2(yq270)*, *pck-2(yq271),* and *got-1.2(yq163)* mutants were obtained by ethyl methanesulfonate (EMS) mutagenesis. The *got-2.2*(*yq341*) mutants were generated by CRISPR/Cas9 system in this laboratory. Integrated arrays, deletion mutants and mutants obtained by EMS mutagenesis were outcrossed with the N2 strain at least four times. All strains used in this study are listed in Supporting information (S1 Table).

### Expression vectors

The expression vectors used in the study were constructed using standard protocols and are listed in Supporting information (S2 Table).

### Ethyl methanesulfonate (EMS) mutagenesis and gene cloning

Synchronized L4-stage *yqIs157* animals expressing MITO-GFP were treated in 50 mM EMS solution for 4 h at 25 °C and then transferred to NGM plates at 20 °C to recover and produce progeny. F2 animals at 24-48 h post-L4 stage were examined for mitochondrial morphology. *yq163*, *yq270* and *yq271* mutants were obtained from a screen of 12,000 haploid genomes. *yq270* and *yq271* mutants showed spherical mitochondria and *yq163* mutants displayed better mitochondrial connection. Mapping of mutations was performed using SNP (single nucleotide polymorphism) mapping and mutation sites were identified with whole genome sequencing.

### Generation of deletion mutants

To generate *got-2.2(yq341)* deletion mutants, a single-guide RNA (sgRNA) targeting sequence (5’-TTTTCTCTACCGCCGTGCG-3’) in the first exon of the *got-2.2* gene was cloned into the Cas9-expressing pDD162 vector. A repair template (5’-TCAACAATGAGCGTTTCCAAGAAGCTTTTCTCTACCGCCGGCTAGCGAAAGTCGTGGTGGTCGCATGTTGAGATGGGACCACCAGA-3’) containing the mutation of interest was designed to remove the cleavage site. An Nhe I restriction site was introduced into the repair template. *dpy-10* was used as a positive control marker. The resulting pDD162 construct (20 ng/μl) containing the sgRNA sequence and repair templates (2 μmol/l) were co-injected with the *dyp-10* sgRNA construct (20 ng/µl) into gonads of young adult animals. Roller F1 worms were singled into new NGM plates, and the F2 animals were examined by PCR amplification (forward primer: 5’-ATCATGCGTTGTACGTACTC-3’; reverse primer: 5’-CAACAATTCCGGCGTACTCC-3’) and restriction enzyme digestion (Nhe I). All mutations were confirmed by sequencing.

### ATP measurements

300 adult animals (48 h post L4) were homogenized in 200 μl of ATP Lite lysis buffer (Vigorous) and subjected to five freeze-thaw cycles in liquid nitrogen. The lysates were boiled for 5 min and then centrifuged at 15,000 ×g for 5 min at 4 °C to collect the supernatants. 20 μl of supernatant was applied to the luciferin-luciferase ATP assay. Luminescence was recorded on a Clarity microplate luminometer (BioTek) between 300 and 600 nm.

### Analysis of body-bending rates

Synchronized adult worms (48 h post L4) were transferred from bacterial lawns to a fresh, unseeded agar plate to remove external bacteria. Individual worms were then transferred to a new unseeded plate containing 1 ml of M9 buffer. Head swings per minute, defined as the return of the head to the original position, were manually counted under a Motic SMZ-168 microscope.

### Measurement of OCR

OCRs were measured at 25 °C using a Seahorse XFe24 analyzer (S7801B). Briefly, 300 adult worms (48 h post L4) were transferred in quadruplicate to a 24-well microplate containing 500 µl of M9 buffer. Basal respiration was measured for a total of 22.5 min with 4.5-min intervals, with each interval consisting of 2 min of mixing, 0.5 min of waiting and 2 min of measuring). Maximal respiration was induced with 20 mM FCCP (carbonyl cyanide 4-(trifluoromethoxy) phenylhydrazone) and detected for a total of 40.5 mins with 4.5-min intervals, with each interval consisting of 2 min of mixing, 0.5 min of waiting and 2 min of measuring.

### Lifespan assays

For each group, 120 synchronized L4 animals were maintained on 4 fresh NGM plates (30 animals per plate) and transferred to a new plate every day until the end of reproduction. Animals that failed to respond to mechanical stimulation were defined as dead. Viable worms were scored every day.

### Body length measurements

Synchronized L4 animals were cultured on fresh NGM plates at 20 °C for 48 h. Worms were then anesthetized with 2.5 mM levamisole in M9 buffer and mounted on 3 % agar pads. Images were acquired using an Axio Imager M2 microscope (Carl Zeiss). Body lengths were quantified from head to tail tip using ZEN software (v.2.3).

### Microscopy and image analysis

*C. elegans* animals were anesthetized in 2.5 mM levamisole in M9 buffer and immobilized on 3% agar pads. Differential interference contrast (DIC) and fluorescence images were captured at 20 °C with an inverted Zeiss LSM 880 confocal microscope equipped with an Alpha Plan-Apochromat 100×/1.46 oil objective. Images were processed using ZEN software (v.2.3).

### TEM analysis

Adult *C. elegans* animals were frozen under high pressure by using a Leica EM ICE system. Samples were subjected to freeze-substitution in a Leica EM AFS2 unit with anhydrous acetone containing 1% osmium tetroxide and 0.1% uranyl acetate. The parameters were: -90 °C for 72 h, -60 °C for 15 h, -30 °C for 15 h, and 0 °C for 5 h. The samples were washed with cold anhydrous acetone, and infiltrated with Embed-812 resin in a graded series (resin:acetone 1:3 for 3 h, 1:1 for 5 h, 3:1 overnight), followed by pure resin for 4 h. The samples were then incubated 60 °C for 48 h for polymerization. Ultrathin sections (70 nm) were cut on a Leica EM UC7 microtome, followed by staining with 2% uranyl acetate for 10 min and lead citrate for 5 min. Images were acquired with a HT7800 (Hitachi) at 80 kV. For TEM analysis of U2OS cells, U2OS cell monolayers were first fixed with 2.5% glutaraldehyde in 0.1 M PBS at 4 °C for 2 h, and then post-fixed with 1% OsO_4_ at 4 °C for 1 h, followed by ethanol dehydration and embedding in EMbed-812. Ultrathin sections (70 nm) were prepared using an UC6 ultramicrotome equipped with a 45° diamond knife (Diatome). The grids were stained with aqueous uranyl acetate and Reynolds lead citrate. Imaging was performed as above.

### Recombinant proteins and GST pull-down

Individual cDNAs were cloned into pET-21a (+) or pGEX-4T-1 and expressed in *E. coli* Rosetta (DE3) strain. His_6_-tagged or GST-fused recombinant proteins were purified with Ni-Sepharose beads (GE Health Care) or Glutathione Sepharose 4B beads (GE Health care), respectively. Proteins were further concentrated and desalted with Amicon Ultra-15 10K centrifugal filters (Merck Millipore), if necessary. For GST pull-down, His_6_-tagged proteins were incubated with GST-fusion proteins immobilized on glutathione Sepharose 4B beads at 4 °C overnight. The beads were washed extensively and bound proteins were resolved by SDS-PAGE and detected with western blotting.

### Western blotting and immunoprecipitation (IP)

For western blotting, adult worms (48 h post-L4 stage) grown on 20 NGM plates (10 cm in diameter) were used to prepare cytosolic and mitochondrial fractions. Cytosolic or mitochondrial samples were separated on 12.5 % SDS-PAGE and analyzed by western blotting. For immunoprecipitation, worms expressing GFP-tagged proteins were lysed in worm lysis buffer (25 mM Tris-HCl, pH 7.5, 50 mM NaCl, 0.1% NP-40, 1 mM PMSF (phenylmethylsulfonyl fluoride) and 1% glycerol, Complete Protease Inhibitor Cocktail (Roche) on ice. The crude lysates were collected by centrifugation at 12,000 g for 10 min at 4 °C. The supernatants were incubated with GFP-Nanoab-Agarose Beads (Lablead) overnight at 4 °C, and washed three times with wash buffer (25 mM Tris-HCl, pH 7.5, 150 mM NaCl, 0.1% NP-40, and 1 mM PMSF). Bound proteins were resolved by SDS-PAGE and detected by western blotting. Antibodies and reagents used in this study are listed in Supporting information (S3 Table).

### RNA interference (RNAi)

Five L4 animals were grown on NGM plates supplemented with 1 mM IPTG (isopropyl β-D-thiogalactoside) and seeded with *E. coli* OP50 expressing double-stranded RNA of genes of interest. The F1 progeny at the L4 stage were transferred to fresh RNAi plates and cultured for 48 h. The animals were then examined for mitochondrial morphology under confocal fluorescence microscope.

### Measurement of OAA levels

Cytosolic and mitochondrial fractions were prepared from *C. elegans* animals and mammalian cells by following previously described methods[52, 53]. Briefly, pellets (approximately 500 μl) of worms synchronized at 48 h post L4 were homogenized in mitochondrial buffer (10 mM MOPS, pH 7.4, 70 mM sucrose, 1 mM EGTA, 210 mM sorbitol) on ice using a Dounce homogenizer. The crude lysates were fractionated by differential centrifugation (450×g, 550×g, and 9,500×g, 10 min each) at 4 °C to pellet the mitochondria. To prepare mitochondria from cultured cells, cell pellets (100 μl) were broken in homogenization buffer (3.5 mM Tris-HCl, pH 7.8, 2.5 mM NaCl, 0.5 mM MgCl_2_, 1 mM PMSF) on ice using a Dounce homogenizer. The lysates were centrifuged sequentially at 1,200× g for 3 min and 15,000× g for 5 min at 4 °C to pellet the mitochondria. Mitochondrial fractions were examined by western blotting using antibodies against HSP-60 and α-Tubulin to indicate mitochondrial and cytosolic fractions, respectively. OAA levels in the mitochondrial and cytosolic fractions were measured by using the Oxaloacetate Assay Kit (Sigma) following the procedures provided by the supplier.

### Treatment with OAA, Asp, Malate or 3-MPA

*C. elegans* embryos were cultured on NGM plates supplemented with OAA (0.1, 1.0, or 5.0 mM), aspartate (5.0 mM), or malate (5.0 mM) and allowed to develop to adults (48 h post L4). The mitochondrial morphology was examined using confocal microscopy. For treatment of mammalian cells, U2OS and HeLa cells were grown to approximately 40% density and then cultured in complete medium supplemented with 3-MPA (0.25, 0.50, or 1.00 mM), OAA (1.0 or 5.0 mM), aspartate (5.0 mM), or malate (5.0 mM). 24 h later, the cells were stained with Mito-CMXRos for 20 min and imaged by confocal microscopy to assess mitochondrial morphology.

### Real-time quantitative PCR (qPCR)

Total RNA was isolated from *C. elegans* at 48 h post L4 stage using TRIzol reagent (Life Technologies). cDNA was synthesized using oligo-(dT18) primers and M-MLV reverse transcriptase (Promega). qPCR was performed on a CFX-96 real-time system (Bio-Rad) with SYBR Premix Ex Taq (Medchem Express). The ΔΔCT method was used for quantification with *act-1* as the reference gene. Oligos used for qPCR are listed in Supporting information (S4 Table).

### Microscale thermophoresis (MST) assays

MST assays were carried out on a Monolith NT.115 system (Nanotemper Technologies). Briefly, 200 nM of target protein (IMMT-1 or CHCH-3) was incubated with varying concentrations of OAA in 20 µl and loaded into NT.115 standard coated capillaries (Nanotemper Technologies). MST measurements were conducted in triplicate at 25 °C with 40 % excitation power and high MST power. Data were processed with NanoTemper software.

### Virtual docking of OAA on CHCH-3

The protein structure file of CHCH-3 was retrieved from the AlphaFold Database (https://alphafold.ebi.ac.uk/) and preprocessed using AutoDockTools 1.5.6[54]. Molecular docking of OAA to CHCH-3 was performed using AutoDock Vina[41, 42]. The resulting ligand-protein poses were analyzed in PyMOL (https://pymol.org/2/) with the most favorable binding pose selected and the hydrogen-bond interactions annotated.

### Liposome assays

Type I lipids (Folch Fraction I, Sigma) were dissolved in chloroform and dried under a nitrogen stream, and further placed under high vacuum (≥3 h) to remove residual solvent. To generate the liposomes, the lipid film was hydrated in a growth buffer (100 mM Tris-HCl, pH7.5, 50 mM NaCl and 50 mM sucrose) at 60 °C for 1 h in the drying oven. Liposomes at 1 mg/ml were incubated with 5 µM of target protein (IMMT-1 and CHCH- 3) and varying concentrations of metabolites (OAA, aspartate, or malate) for 30 min at 20 °C. 5-7 µl of liposome suspension was loaded on a grid coated with formvar and incubated for 1 min. After removal of excess liposome solution, the grid was stained with 2 % uranyl acetate solution and imaged with an HT7800 (Hitachi) transmission electron microscope operating at 80 kV.

### Statistics and reproducibility

Statistical analyses were performed using GraphPad Software (version 8.2.1, Dotmatics) and IBM SPSS statistics (version 21.0). Data are presented as mean ± s.e.m (standard error of the mean). Comparisons between two groups were made by using the two-tailed unpaired Student’s *t*-test. Comparisons of multiple groups were assessed by using one-way or two-way analysis of variance (ANOVA) followed by Dunnett’s, Sidak’s, or Tukey’s post-hoc test. ***, *P*<0.001; **, *P* < 0.01; *, *P* < 0.05; ns, not significant (*P* > 0.05).

## Acknowledgments

We thank I. Hanson for proofreading the manuscript and the Caenorhabditis Genetics Center for *C. elegans* strains used in this study.

## Author Contributions

**Conceptualization:** Jie Zhang, Chonglin Yang

**Data curation:** Jie Zhang, Qian Shan

**Formal analysis:** Jie Zhang, Qian Shan

**Funding acquisition:** Jie Zhang, Chonglin Yang

**Investigation:** Jie Zhang, Qian Shan, Xin Wang, Meijiao Li, Yang Yang, Mei Duan, Ruofeng Tang, Junxiang Zhou, Fengyang Wang

**Methodology:** Jie Zhang, Qian Shan, Xin Wang, Yuehui Shi, Kai Jiang

**Project administration:** Jie Zhang, Chonglin Yang

**Resources:** Chonglin Yang

**Supervision:** Jie Zhang, Chonglin Yang

**Writing – original draft:** Jie Zhang, Chonglin Yang

**Writing – review & editing:** Jie Zhang, Qian Shan, Chonglin Yang

## Supporting information

**S1 Fig.**
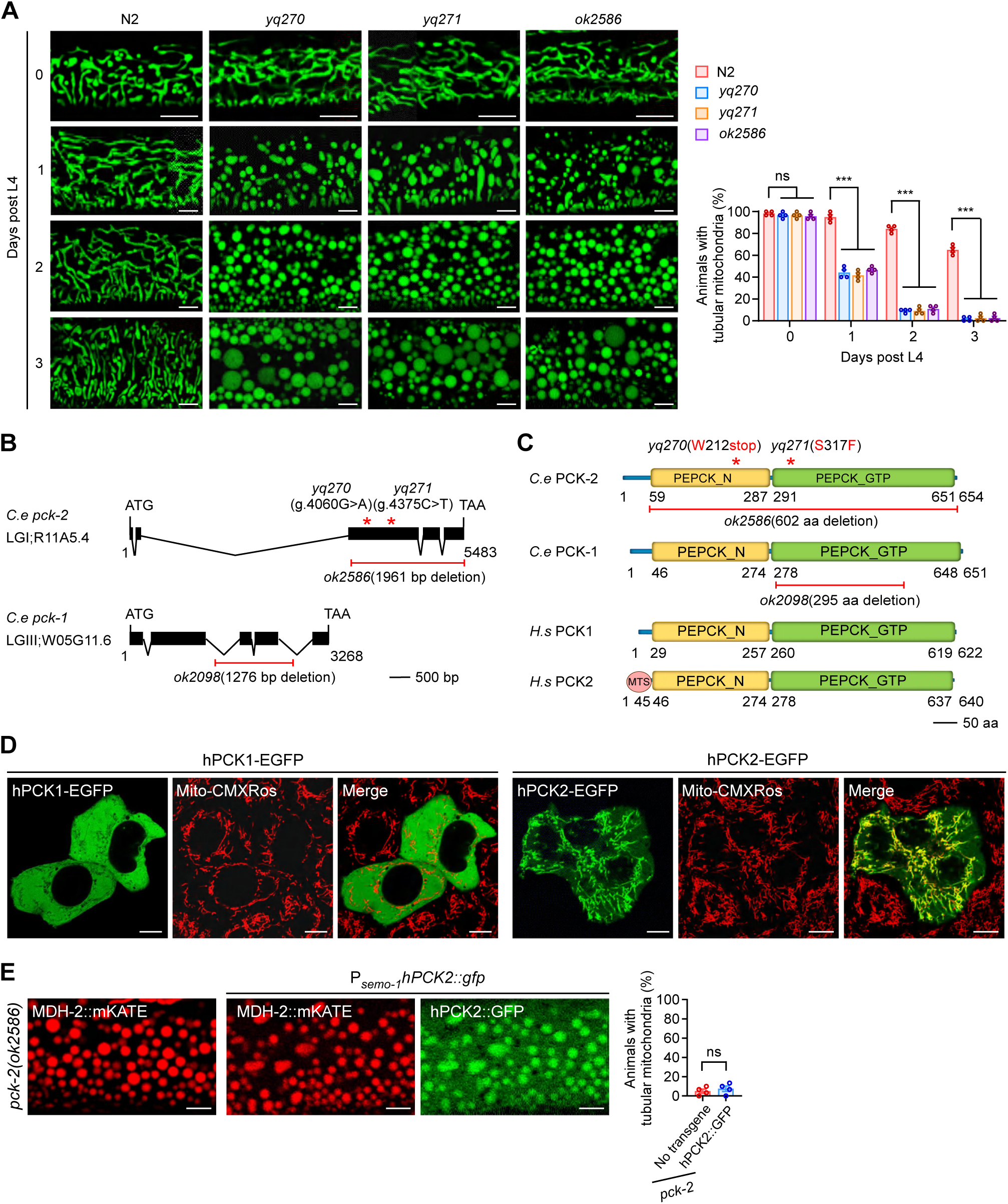
Mutations in *pck-2* cause abnormal mitochondrial morphology in *C. elegans*. (**A**) Images (left) and quantification (right) of mitochondria in the hypodermis of N2 (wild type), *yq270*, *yq271* and *ok2586* animals. Adult animals were examined at the indicated ages after the 4^th^ larval (L4) stage. A total of 120 animals were examined in *n*= 4 repeats for each genotype. Bars, 5 μm. **(B)** Schematic depiction of *C. elegans pck-2* and *pck-1* genes. Filled boxes and thin lines represent exons and introns, respectively. The locations of the *yq270* and *yq271* point mutations are indicated with red asterisks. *ok2586* and *ok2098* deletions are indicated with red horizontal lines. **(C)** Diagram of *C. elegans* (*C.e.*) PCK-2, *C.e.* PCK-1, human (*Homo sapiens*, *H.s.*) PCK1 and *H.s* PCK2 proteins. Mutation sites and deletion regions are indicated in *C.e.* PCK-2 and *C.e.* PCK-1. The mitochondrion-targeting sequence (MTS) of PCK2 is indicated in the red circle. Yellow and green boxes indicate PEPCK_N and PEPCK_GTP domains, respectively. **(D)** Representative images of mitochondria in HeLa cells with transient expression of human PCK1-EGFP and human PCK2-EGFP. Mitochondria were stained with Mito-CMXRos. Bars, 10 μm. **(E)** Full-length hPCK2 fails to ameliorate the abnormal mitochondria in *C. elegans pck-2(ok2586)* mutants. hPCK2::GFP driven by the *semo-1* promoter (P*_semo-1_hPCK2::gfp*) was expressed in hypodermis of *C. elegans pck-2(ok2586)* mutants expressing MDH-2::mKATE. Left, representative images of mitochondria; right, quantification. Bars, 5 μm. A total of 120 animals were examined for each group in *n*= 4 independent repeats. For quantifications, data points represent (mean ± s.e.m.). *P* values were determined using one-way ANOVA in (A) or the two-tailed unpaired Student’s *t*-test in (E). \*\*\**P* < 0.001; \*\**P* < 0.01; \**P* < 0.05; ns, not significant (*P* > 0.05).

**S2 Fig.**
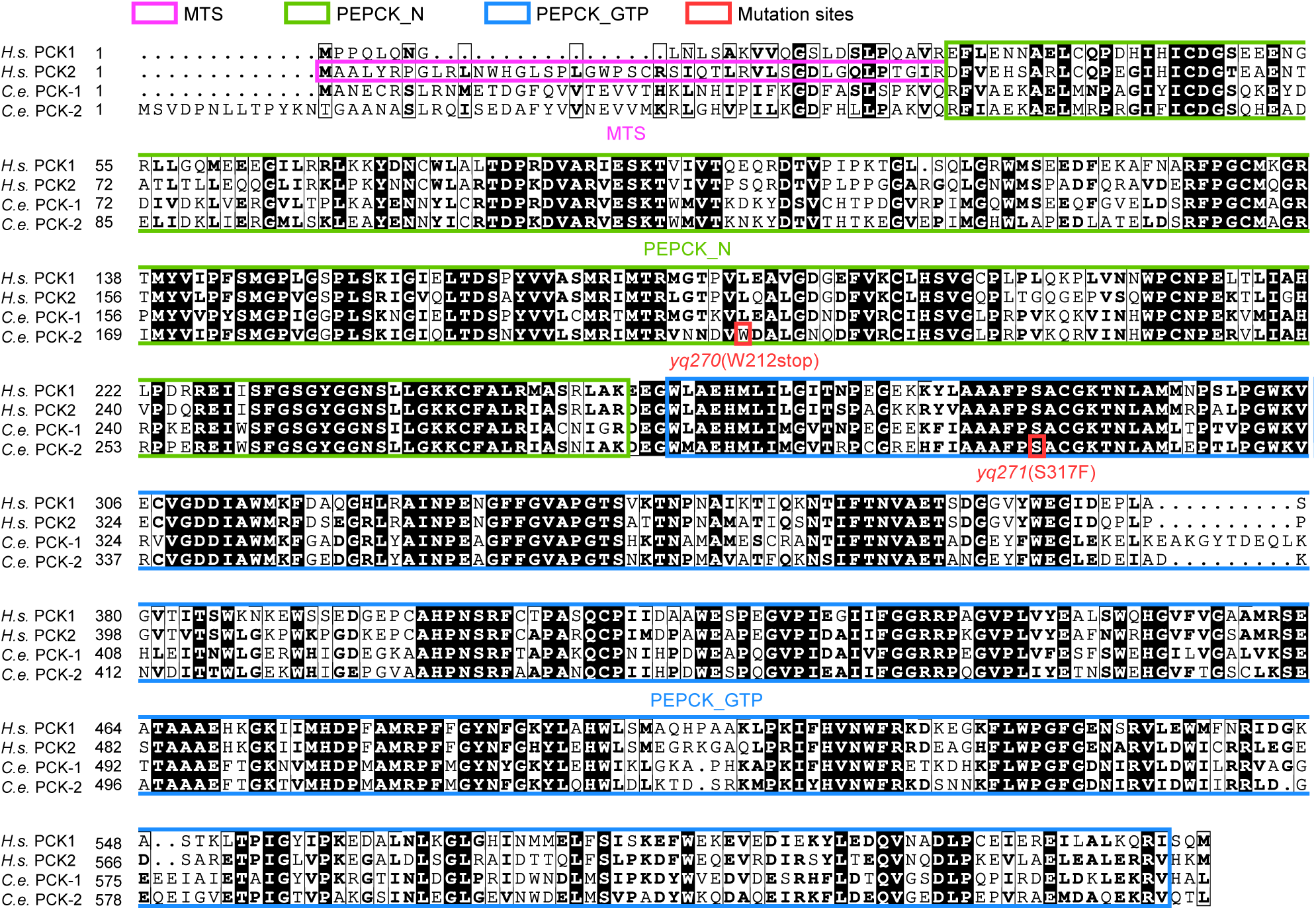
Amino acid sequence alignment of human (*H.s.*) PCK1 and PCK2 with *C. elegans* PCK-1 and PCK-2. The mitochondrion-targeting sequence (MTS) of *H.s.* PCK2 is indicated in the violet box. The PEPCK_N and PEPCK_GTP domains are indicated in green and blue boxes, respectively. The residues affected by the *yq270* and *yq271* point mutations in *C.e.* PCK-2 are labeled in red.

**S3 Fig.**
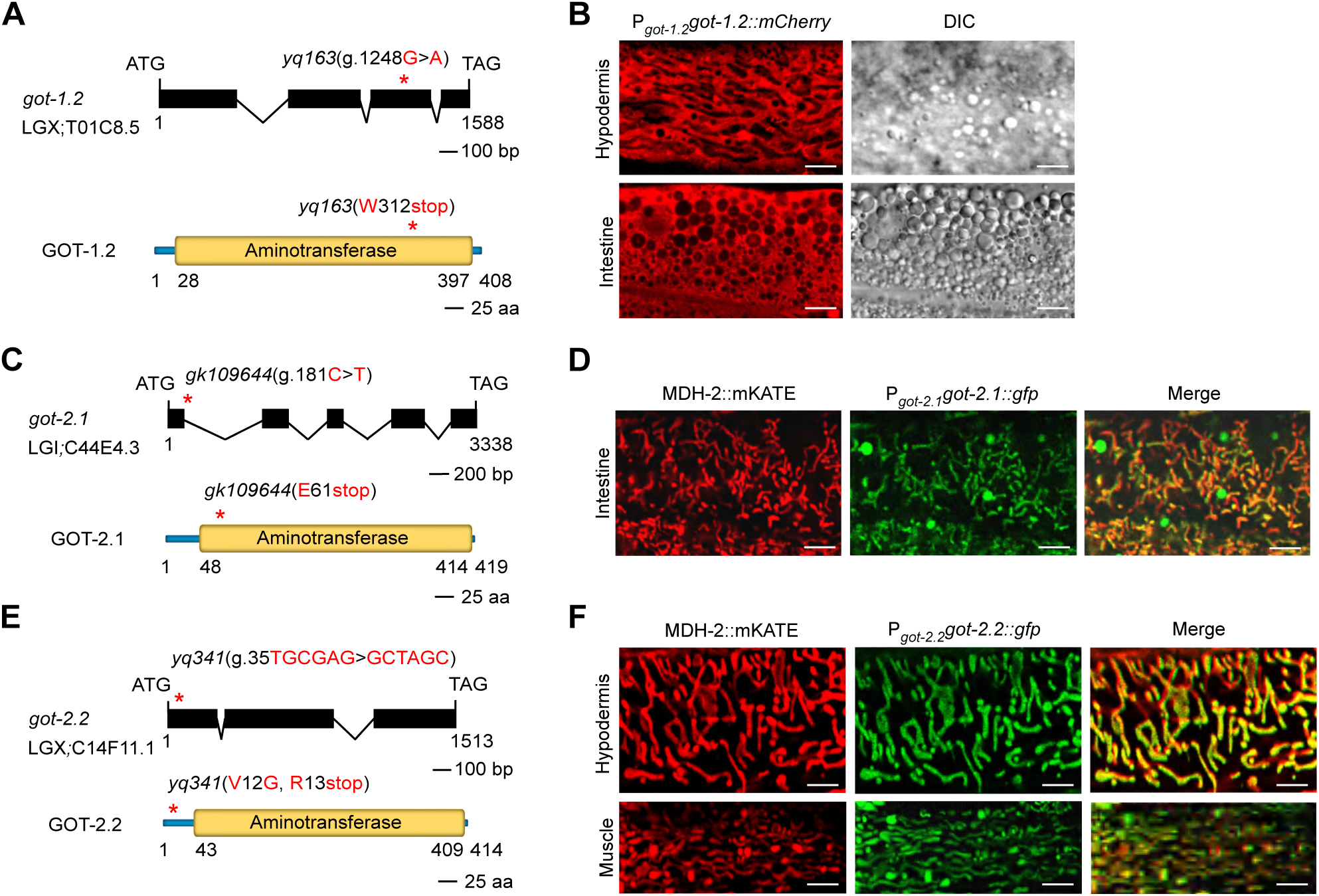
Characterization of glutamate-oxaloacetate transaminases. **(A)** Diagram of the *C. elegans got-1.2* gene (top) and the encoded protein (bottom). In the gene, filled boxes and thin lines represent exons and introns, respectively. The location of the *yq163* point mutation is indicated with the red asterisk. In the protein, the yellow box indicates the aminotransferase domain. **(B)** Expression pattern of GOT-1.2::mCh driven by the *got-1.2* promoter (P*_got-1.2_got-1.2::mcherry*) in adult N2 hypodermal and intestinal cells. DIC (differential interference contrast) and fluorescence images are shown for each tissue. Bars, 5 µm. **(C)** Diagram of the *C. elegans got-2.1* gene (top) and the encoded protein (bottom). The *gk109644* point mutation is indicated with the red asterisk. The yellow box indicates the aminotransferase domain. **(D)** Expression pattern of GOT-2.1::GFP under the control of the *got-2.1* promoter (P*_got-2.1_got-2.1::gfp*) in adult N2 intestine cells. Mitochondria were labeled with MDH-2::mKATE. Bars, 5 µm. **(E)** Diagram of the *C. elegans got-2.2* gene (top) and the encoded protein (bottom). The *yq341* point mutation is indicated with the red asterisk. The yellow box indicates the aminotransferase domain. **(F)** Expression pattern of GOT-2.2::GFP driven by the *got-2.2* promoter (P*_got-2.2_got-2.2::gfp*) in adult N2 hypodermal and muscle cells. Mitochondria were labeled with MDH-2::mKATE. Bars, 5 µm.

**S4 Fig.**
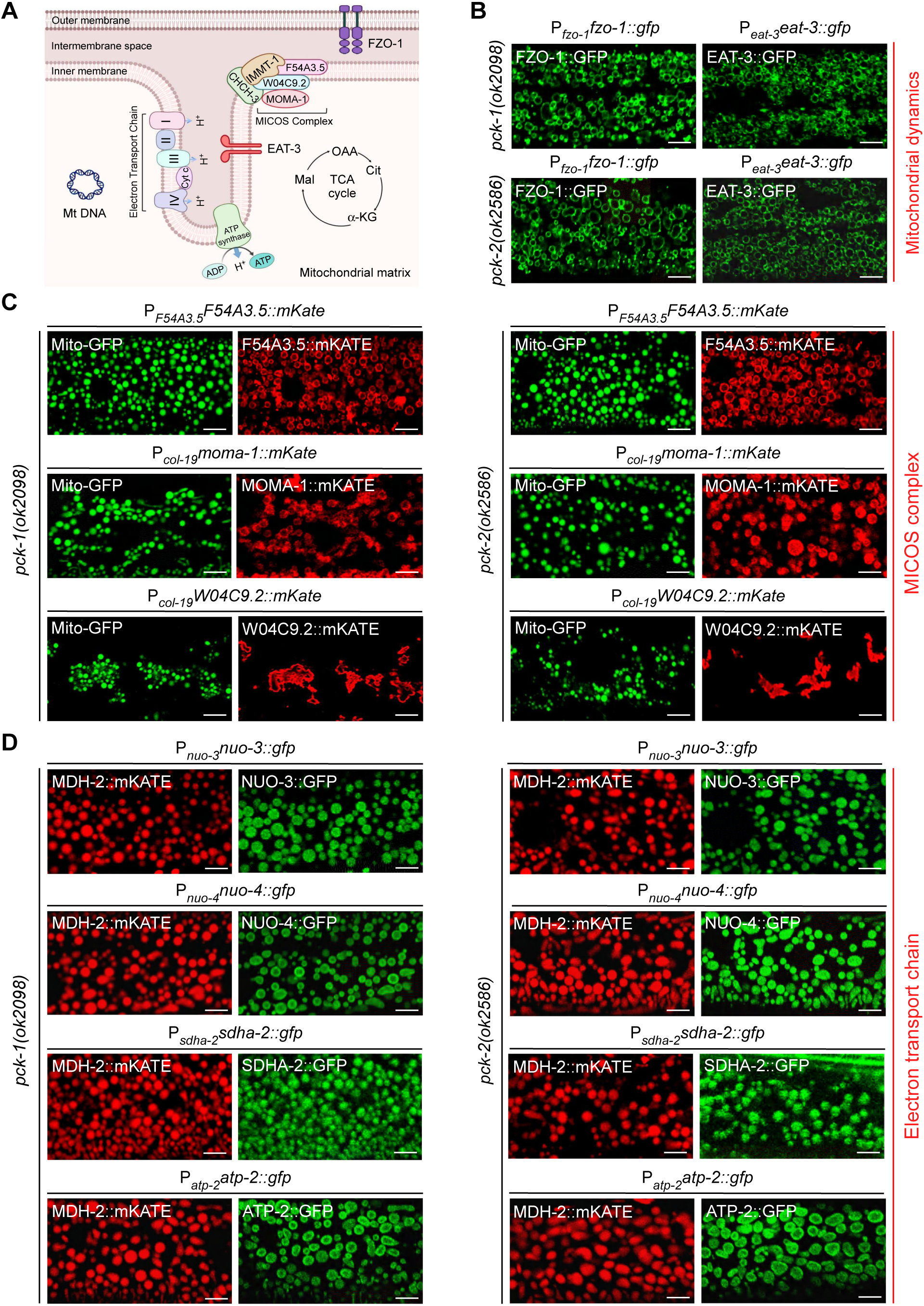
The effects of overexpression of key mitochondrial proteins on mitochondrial defects in *pck-1* and *pck-2* mutants. **(A)** Schematic depiction of key regulators of mitochondrial fusion and cristae formation and maintenance. **(B)** Representative images of mitochondria in the hypodermis in *pck-1*(*ok2098*) and *pck-2*(*ok2586*) mutants overexpressing FZO-1::GFP (P*_fzo-1_fzo-1::gfp*) or EAT-3::GFP (P*_eat-3_eat-3::gfp*). Bars, 5 μm. **(C)** Representative images of mitochondria (labeled with MITO-GFP) in the hypodermis in *pck-1*(*ok2098*) and *pck-2*(*ok2586*) mutants overexpressing F54A3.5::mKATE (P*_F54A3.5_F54A3.5::mKate*), MOMA-1::mKATE (P*_moma-1_moma-1::mKate*) or W04C9.2::mKATE (P*_col-19_W04C9.2::mKate*). Bars, 5 μm. **(D)** Representative images of mitochondria (labeled with MDH-2::mKATE) in the hypodermis in *pck-1*(*ok2098*) and *pck-2*(*ok2586*) mutants overexpressing NUO-3::GFP (P*_nuo-3_nuo-3:gfp*), NUO-4::GFP (P*_nuo-4_nuo-4:: gfp*), SDHA-2::GFP (P*_sdha-2_sdha-2:: gfp*) or ATP-2::GFP (P*_atp-2_atp-2:: gfp*). Bars, 5 μm.

**S5 Fig.**
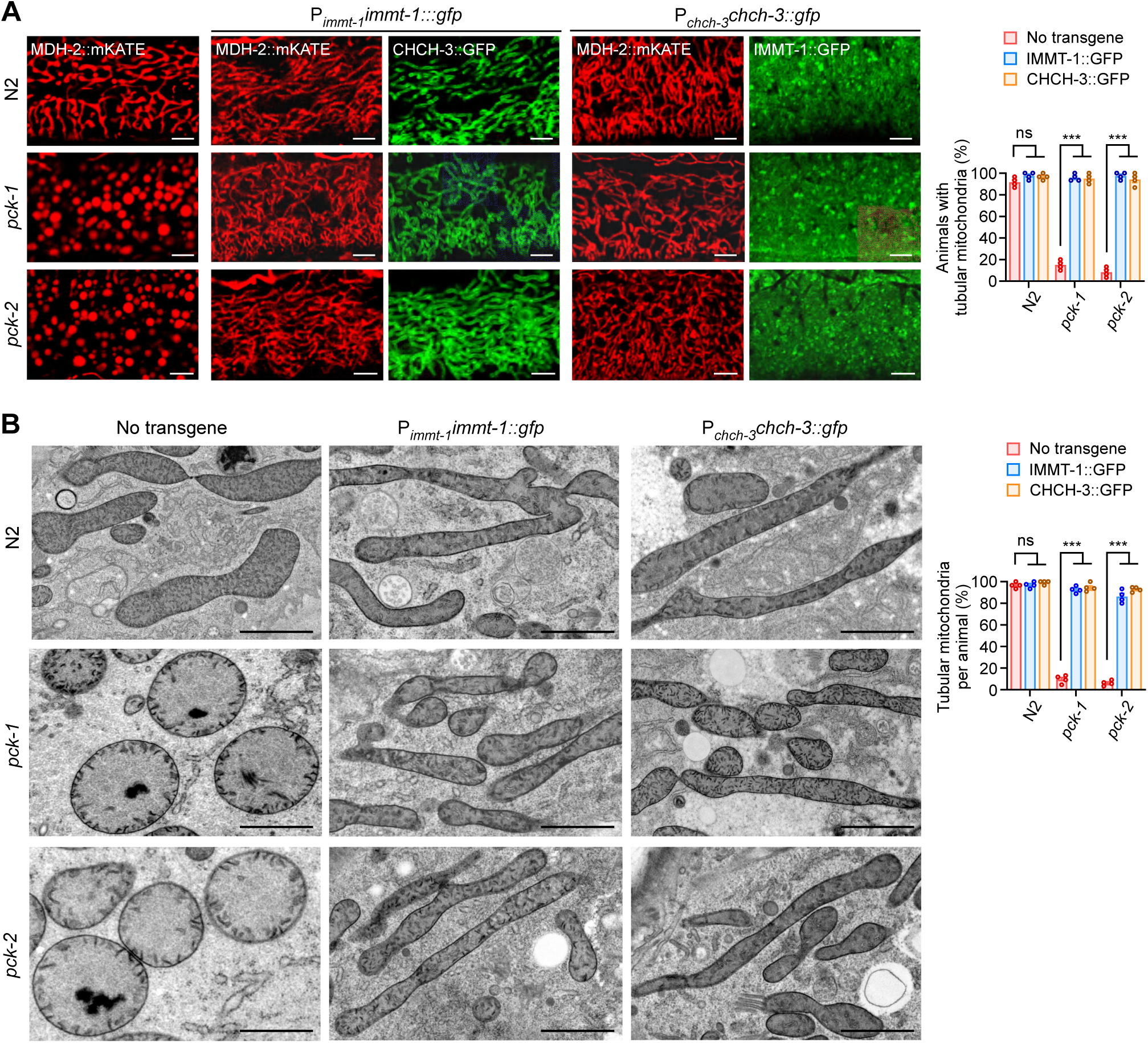
Reinforced expression of IMMT-1 and CHCH-3 restores mitochondrial structure in *pck-1* and *pck-2* single mutants. **(A)** Representative images (left) and quantification (right) of mitochondria (labeled with MDH-2::mKATE) in the hypodermis of N2, *pck-1*(*ok2098*) single mutants, and *pck-2*(*ok2586*) single mutants without or with overexpression of IMMT-1::GFP (P*_immt-1_immt-1::gfp*) or CHCH-3::GFP (P*_chch-3_chch-3::gfp*). Bars, 5 μm. A total of 120 animals were examined for each genotype in *n*= 4 repeats. **(B)** TEM images (left) and quantification (right) of mitochondria in the hypodermis of N2, *pck-1*(*ok2098*) single mutants, and *pck-2*(*ok2586*) single mutants without or with overexpression of IMMT-1::GFP (P*_immt-1_immt-1::gfp*) or CHCH-3::GFP (P*_chch-3_chch-3::gfp*). A total of 400 mitochondria from *n*= 4 animals were examined for each genotype. Bars, 1 μm. For quantifications, data points represent (mean ± s.e.m.). *P* values were determined using one-way ANOVA in (A and B). \*\*\**P* < 0.001; \*\**P* < 0.01; \**P* < 0.05; ns, not significant (*P* > 0.05).

**S6 Fig.**
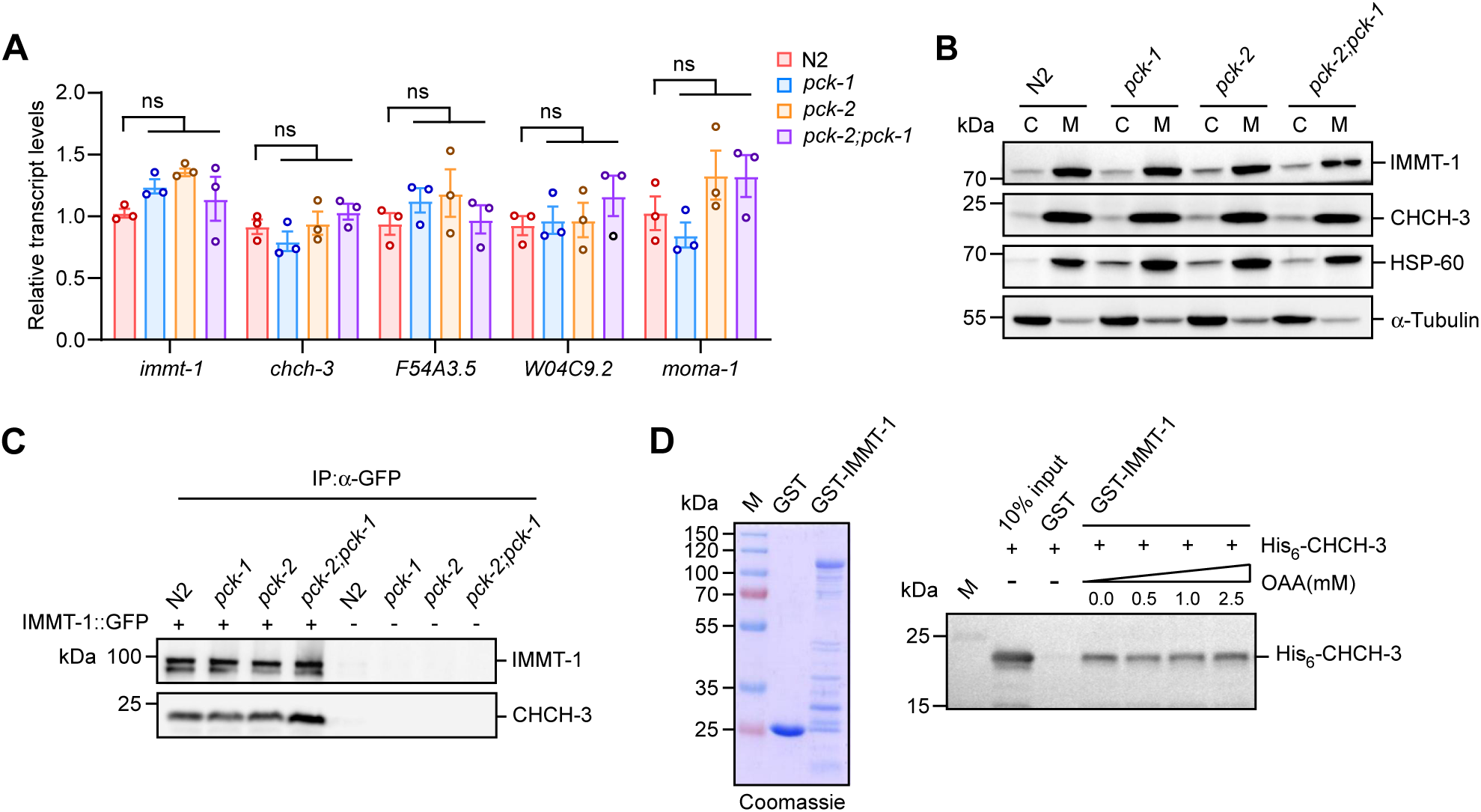
Analysis of the effects of OAA on expression of MICOS subunits and the interaction of IMMT-1 with CHCH-3. **(A)** Relative levels of mRNAs encoding the indicated MICOS subunits. *n*=3 independent experiments. **(B)** Western blotting of IMMT-1 and CHCH-3 in purified mitochondria from animals of the indicated genotypes. IMMT-1, CHCH-3, HSP-60 (mitochondrial marker) and α-tubulin (cytosolic marker) were detected using the corresponding antibodies. **(C)** Co-IP of endogenous CHCH-3 with IMMT-1::GFP (knock-in) from animals with the indicated genotypes. IPs were performed with α-GFP antibody and precipitated proteins were detected with IMMT-1 and CHCH-3 antibodies. **(D)** His_6_-CHCH-3 is pulled down by GST-IMMT-1 in the presence of increasing concentrations of OAA. His_6_-CHCH-3 was detected with His_6_ antibody. Coomassie blue staining of purified GST and GST-IMMT-1 is shown on the left. For quantifications, data points represent (mean ± s.e.m.). *P* values were determined using one-way ANOVA in (A). \*\*\**P* < 0.001; \*\**P* < 0.01; \**P* < 0.05; ns, not significant (*P* > 0.05).

**S7 Fig.**
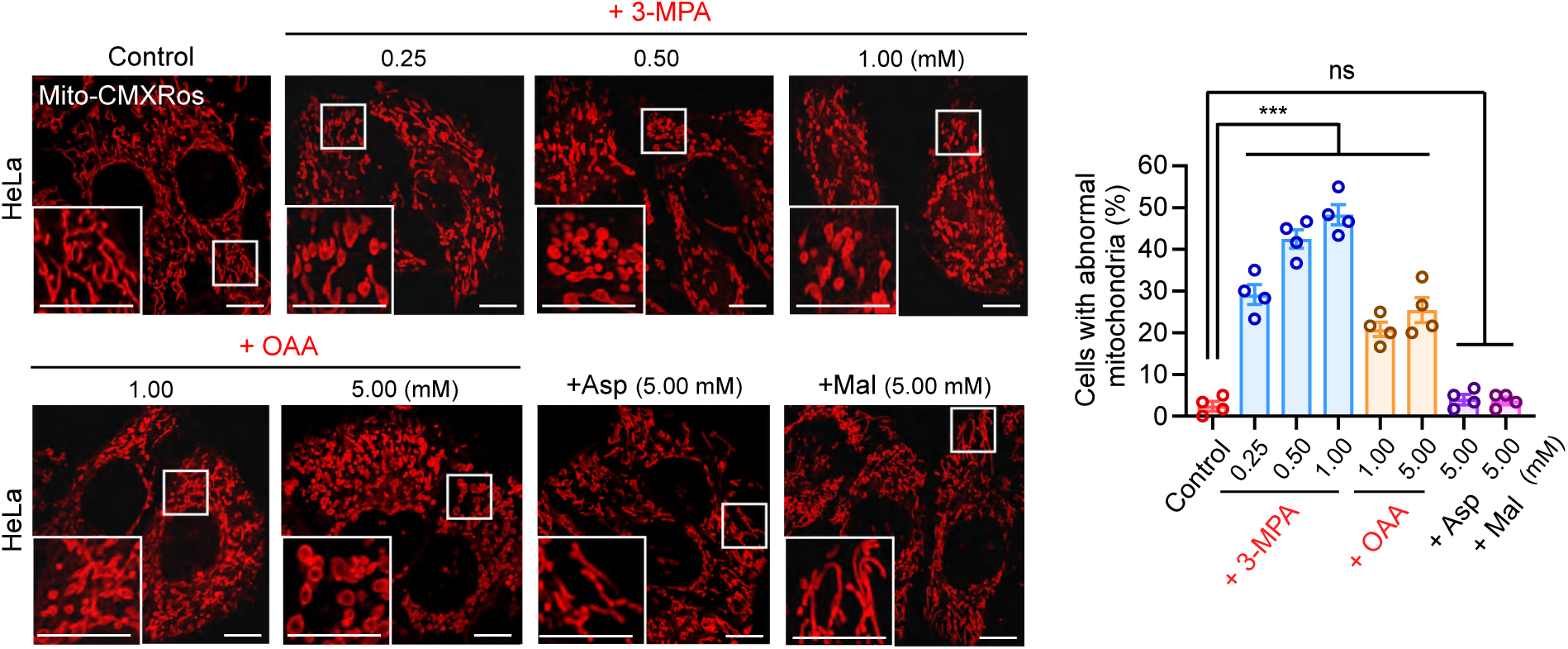
Inhibition of PCK or addition of exogenous OAA leads to abnormal mitochondria in HeLa cells. Representative images (left) and quantification (right) of abnormal mitochondria in HeLa cells treated with 3-MPA (inhibitor of PCK1 and PCK2), OAA, Asp, and Mal at the indicated concentrations. Mitochondria were labeled with Mito-CMXRos. Boxed regions are magnified (2.5×) in the bottom left. Bars, 10 μm. A total of ≥240 cells were scored in each treatment in 4 repeats. For quantifications, data points represent (mean ± s.e.m.). *P* values were determined using one-way ANOVA. \*\*\**P* < 0.001; \*\**P* < 0.01; \**P* < 0.05; ns, not significant (*P* > 0.05).

**S1 Table.**
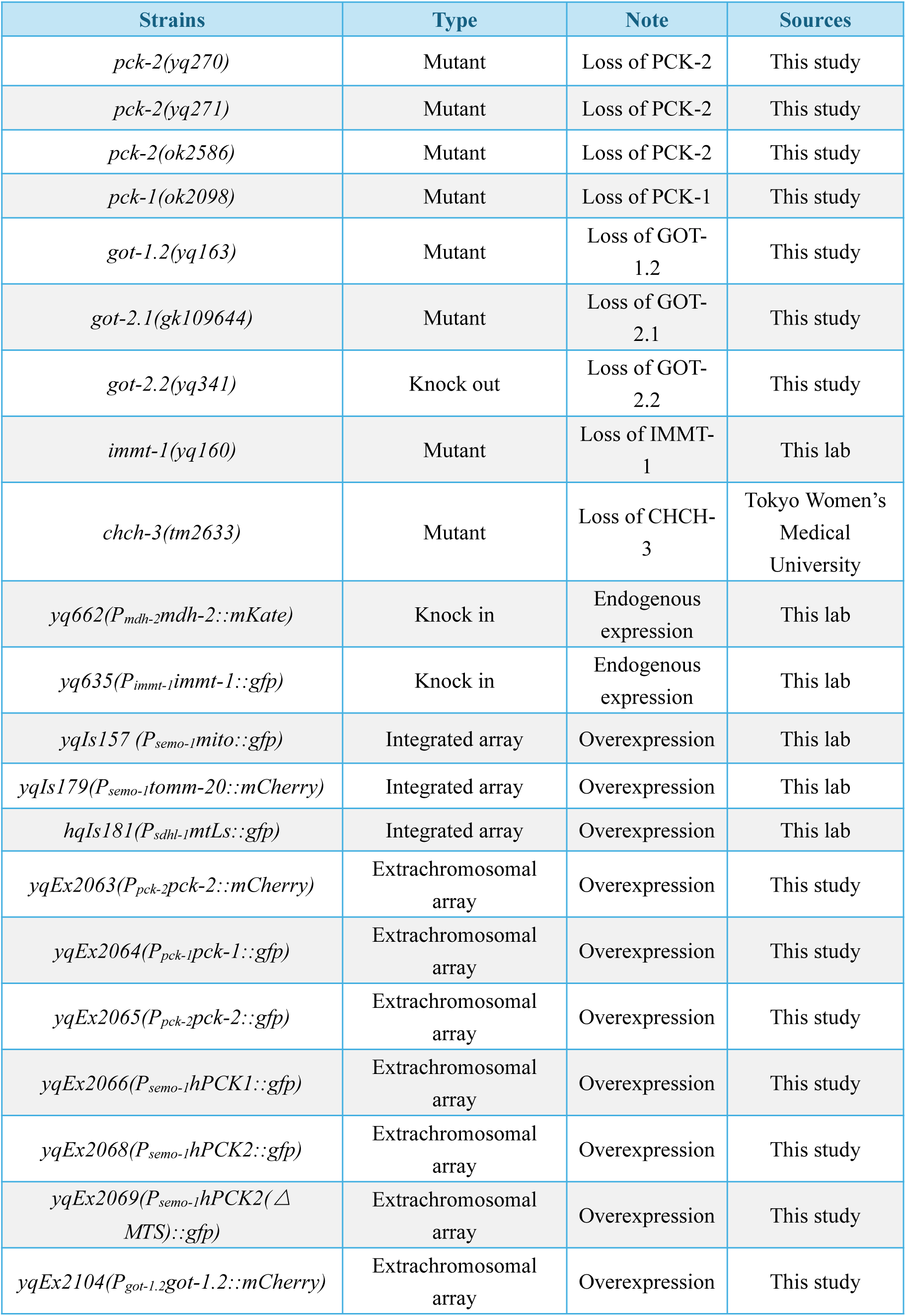
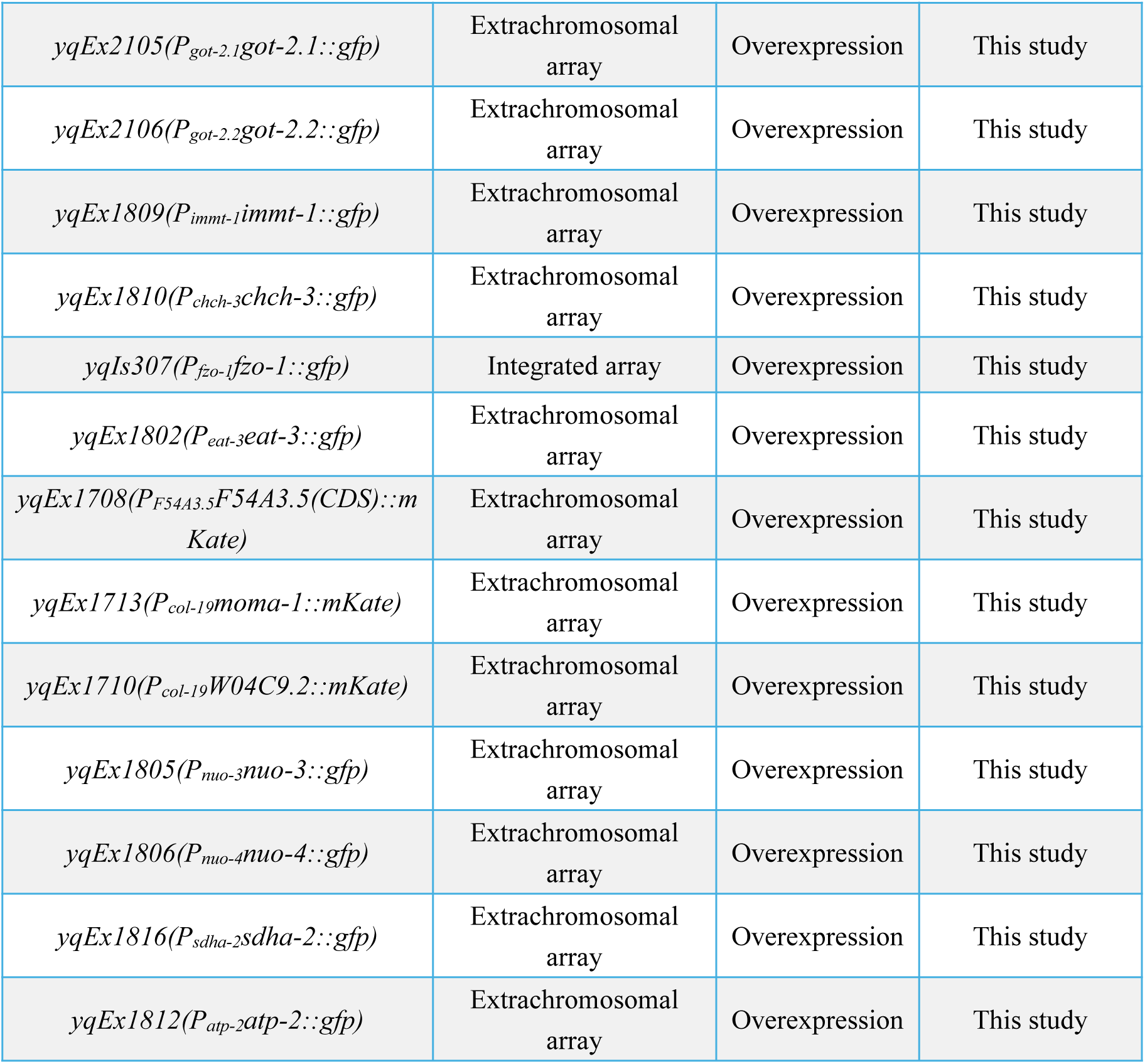
*Caenorhabditis elegans* strains used in this study.

**S2 Table.**
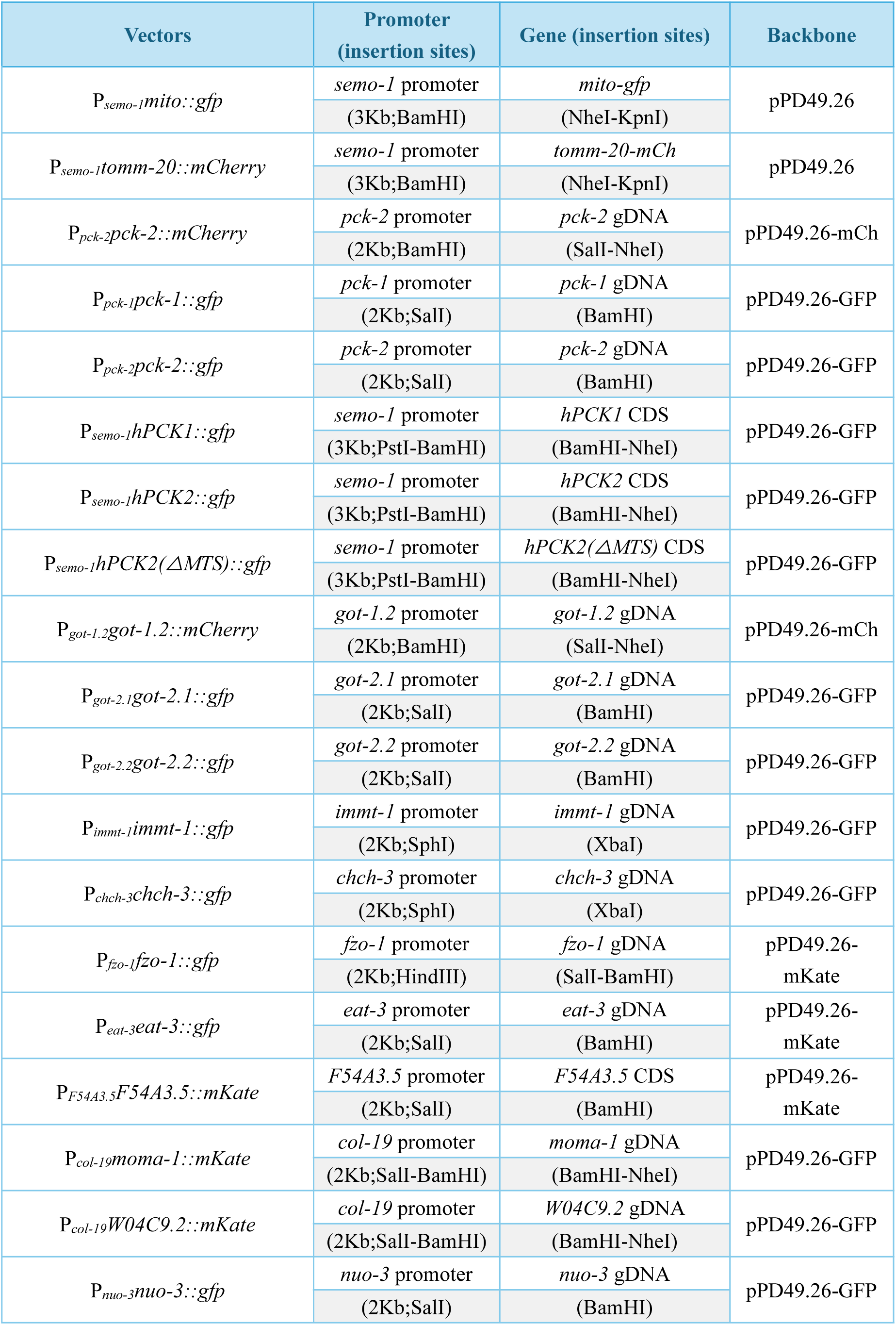
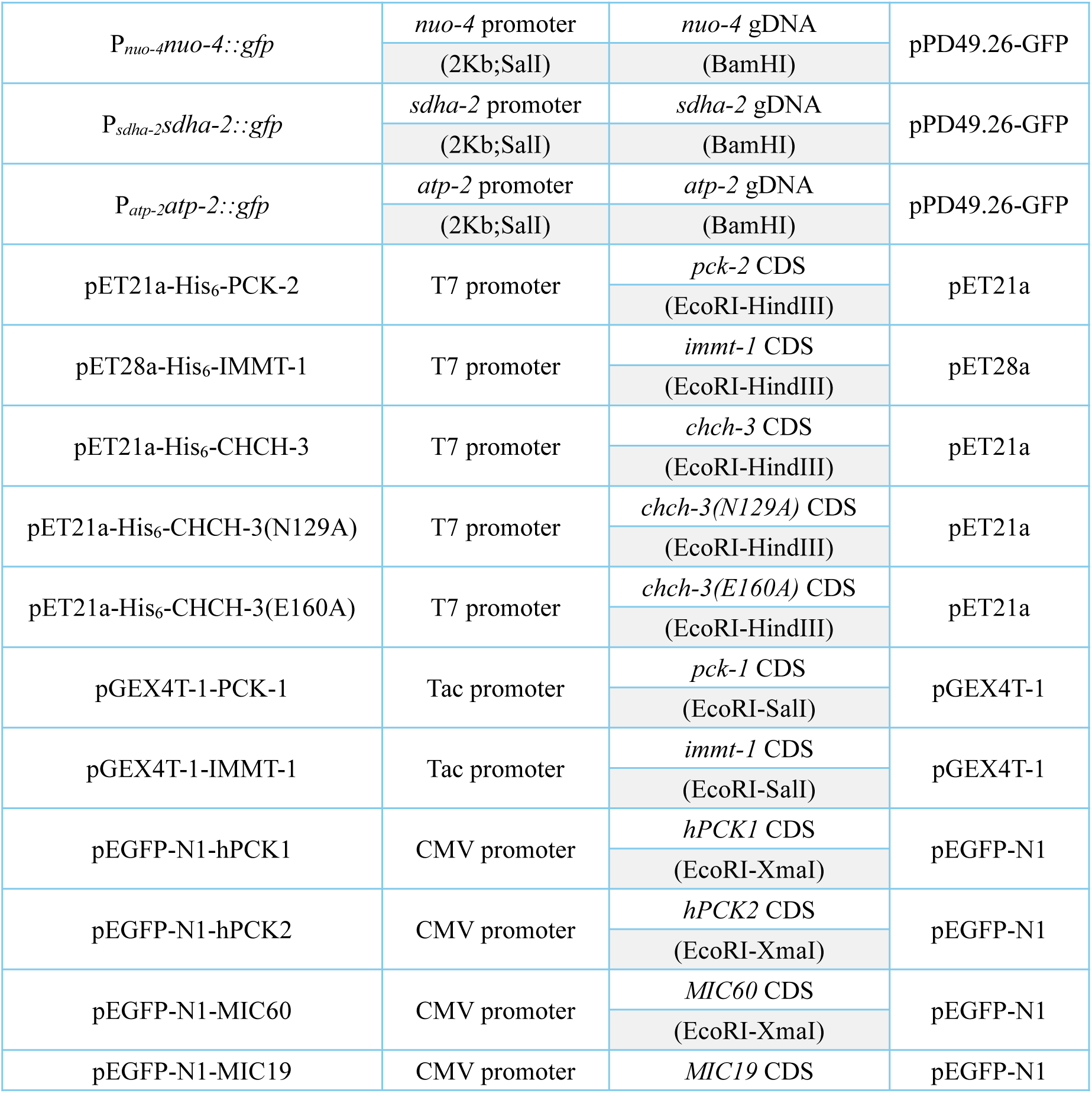
Expression vectors.

| Vectors | Promoter<br>(insertion sites) | Gene (insertion sites) | Backbone |
| --- | --- | --- | --- |
| $P_{semo-1mito::gfp}$ | *semo-1* promoter | *mito-gfp* | pPD49.26 |
|  | (3Kb;BamHI) | (NheI-KpnI) |  |
| $P_{semo-1tomm-20::mCherry}$ | *semo-1* promoter | *tomm-20-mCh* | pPD49.26 |
|  | (3Kb;BamHI) | (NheI-KpnI) |  |
| $P_{pck-2pck-2::mCherry}$ | *pck-2* promoter | *pck-2* gDNA | pPD49.26-mCh |
|  | (2Kb;BamHI) | (Sall-NheI) |  |
| $P_{pck-1pck-1::gfp}$ | *pck-1* promoter | *pck-1* gDNA | pPD49.26-GFP |
|  | (2Kb;Sall) | (BamHI) |  |
| $P_{pck-2pck-2::gfp}$ | *pck-2* promoter | *pck-2* gDNA | pPD49.26-GFP |
|  | (2Kb;Sall) | (BamHI) |  |
| $P_{semo-1hPCK1::gfp}$ | *semo-1* promoter | *hPCK1* CDS | pPD49.26-GFP |
|  | (3Kb;PstI-BamHI) | (BamHI-NheI) |  |
| $P_{semo-1hPCK2::gfp}$ | *semo-1* promoter | *hPCK2* CDS | pPD49.26-GFP |
|  | (3Kb;PstI-BamHI) | (BamHI-NheI) |  |
| $P_{semo-1hPCK2(\Delta MTS)::gfp}$ | *semo-1* promoter | *hPCK2(ΔMTS)* CDS | pPD49.26-GFP |
|  | (3Kb;PstI-BamHI) | (BamHI-NheI) |  |
| $P_{got-1.2got-1.2::mCherry}$ | *got-1.2* promoter | *got-1.2* gDNA | pPD49.26-mCh |
|  | (2Kb;BamHI) | (Sall-NheI) |  |
| $P_{got-2.1got-2.1::gfp}$ | *got-2.1* promoter | *got-2.1* gDNA | pPD49.26-GFP |
|  | (2Kb;Sall) | (BamHI) |  |
| $P_{got-2.2got-2.2::gfp}$ | *got-2.2* promoter | *got-2.2* gDNA | pPD49.26-GFP |
|  | (2Kb;Sall) | (BamHI) |  |
| $P_{immt-1immt-1::gfp}$ | *immt-1* promoter | *immt-1* gDNA | pPD49.26-GFP |
|  | (2Kb;SphI) | (XbaI) |  |
| $P_{chch-3chch-3::gfp}$ | *chch-3* promoter | *chch-3* gDNA | pPD49.26-GFP |
|  | (2Kb;SphI) | (XbaI) |  |
| $P_{fzo-1fzo-1::gfp}$ | *fzo-1* promoter | *fzo-1* gDNA | pPD49.26-mKate |
|  | (2Kb;HindIII) | (Sall-BamHI) |  |
| $P_{eat-3eat-3::gfp}$ | *eat-3* promoter | *eat-3* gDNA | pPD49.26-mKate |
|  | (2Kb;Sall) | (BamHI) |  |
| $P_{F54A3.5F54A3.5::mKate}$ | *F54A3.5* promoter | *F54A3.5* CDS | pPD49.26-mKate |
|  | (2Kb;Sall) | (BamHI) |  |
| $P_{col-19moma-1::mKate}$ | *col-19* promoter | *moma-1* gDNA | pPD49.26-GFP |
|  | (2Kb;Sall-BamHI) | (BamHI-NheI) |  |
| $P_{col-19W04C9.2::mKate}$ | *col-19* promoter | *W04C9.2* gDNA | pPD49.26-GFP |
|  | (2Kb;Sall-BamHI) | (BamHI-NheI) |  |
| $P_{nuo-3nuo-3::gfp}$ | *nuo-3* promoter | *nuo-3* gDNA | pPD49.26-GFP |
|  | (2Kb;Sall) | (BamHI) |  |
| $P_{nuo-4nuo-4::gfp}$ | *nuo-4* promoter | *nuo-4* gDNA | pPD49.26-GFP |
|  | (2Kb;Sall) | (BamHI) |  |
| $P_{sdha-2sdha-2::gfp}$ | *sdha-2* promoter | *sdha-2* gDNA | pPD49.26-GFP |
|  | (2Kb;Sall) | (BamHI) |  |
| $P_{atp-2atp-2::gfp}$ | *atp-2* promoter | *atp-2* gDNA | pPD49.26-GFP |
|  | (2Kb;Sall) | (BamHI) |  |
| pET21a-His<sub>6</sub>-PCK-2 | T7 promoter | *pck-2* CDS | pET21a |
|  |  | (EcoRI-HindIII) |  |
| pET28a-His<sub>6</sub>-IMMT-1 | T7 promoter | *immt-1* CDS | pET28a |
|  |  | (EcoRI-HindIII) |  |
| pET21a-His<sub>6</sub>-CHCH-3 | T7 promoter | *chch-3* CDS | pET21a |
|  |  | (EcoRI-HindIII) |  |
| pET21a-His<sub>6</sub>-CHCH-3(N129A) | T7 promoter | *chch-3(N129A)* CDS | pET21a |
|  |  | (EcoRI-HindIII) |  |
| pET21a-His<sub>6</sub>-CHCH-3(E160A) | T7 promoter | *chch-3(E160A)* CDS | pET21a |
|  |  | (EcoRI-HindIII) |  |
| pGEX4T-1-PCK-1 | Tac promoter | *pck-1* CDS | pGEX4T-1 |
|  |  | (EcoRI-Sall) |  |
| pGEX4T-1-IMMT-1 | Tac promoter | *immt-1* CDS | pGEX4T-1 |
|  |  | (EcoRI-Sall) |  |
| pEGFP-N1-hPCK1 | CMV promoter | *hPCK1* CDS | pEGFP-N1 |
|  |  | (EcoRI-XmaI) |  |
| pEGFP-N1-hPCK2 | CMV promoter | *hPCK2* CDS | pEGFP-N1 |
|  |  | (EcoRI-XmaI) |  |
| pEGFP-N1-MIC60 | CMV promoter | *MIC60* CDS | pEGFP-N1 |
|  |  | (EcoRI-XmaI) |  |
| pEGFP-N1-MIC19 | CMV promoter | *MIC19* CDS | pEGFP-N1 |

**S3 Table.**
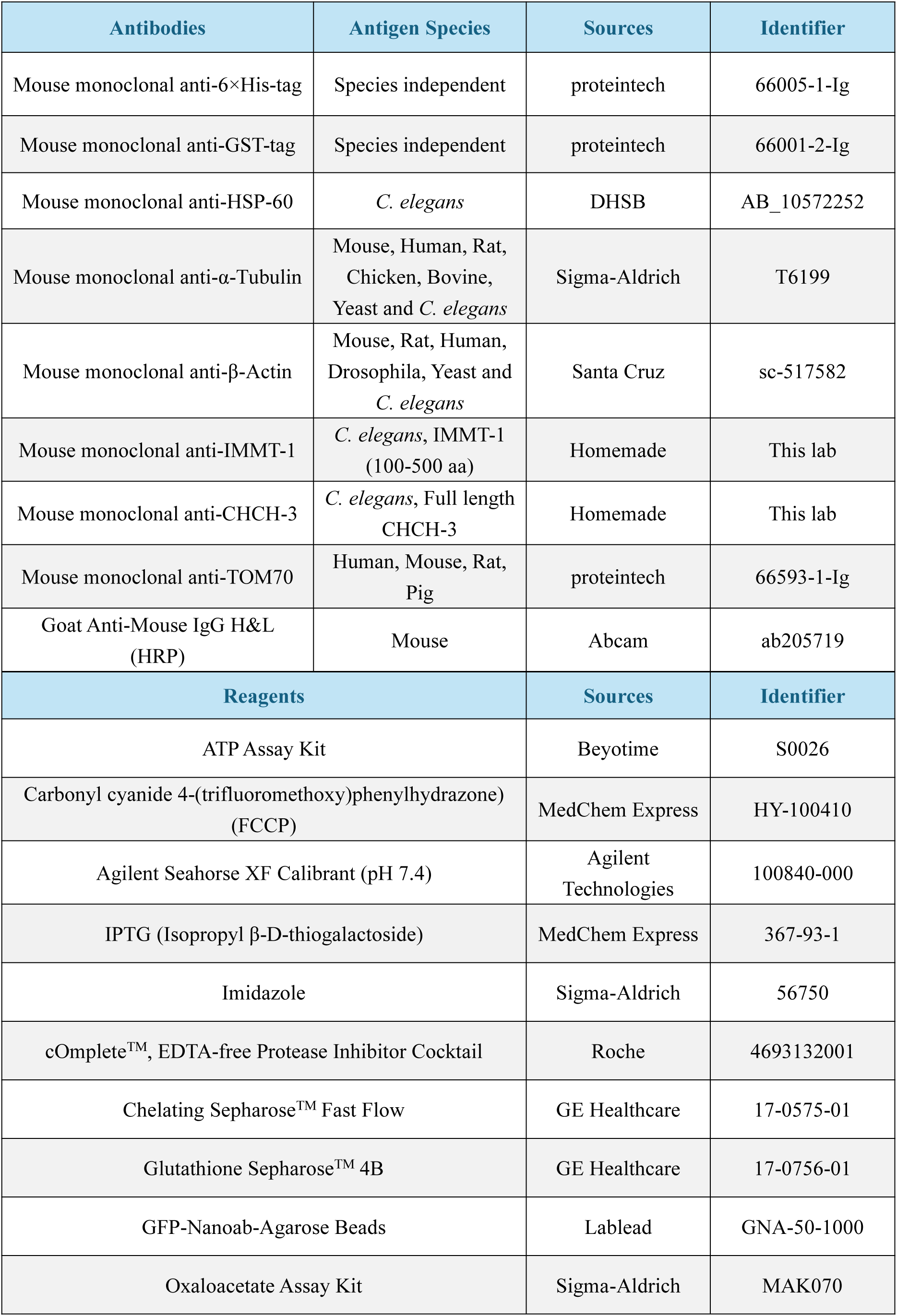
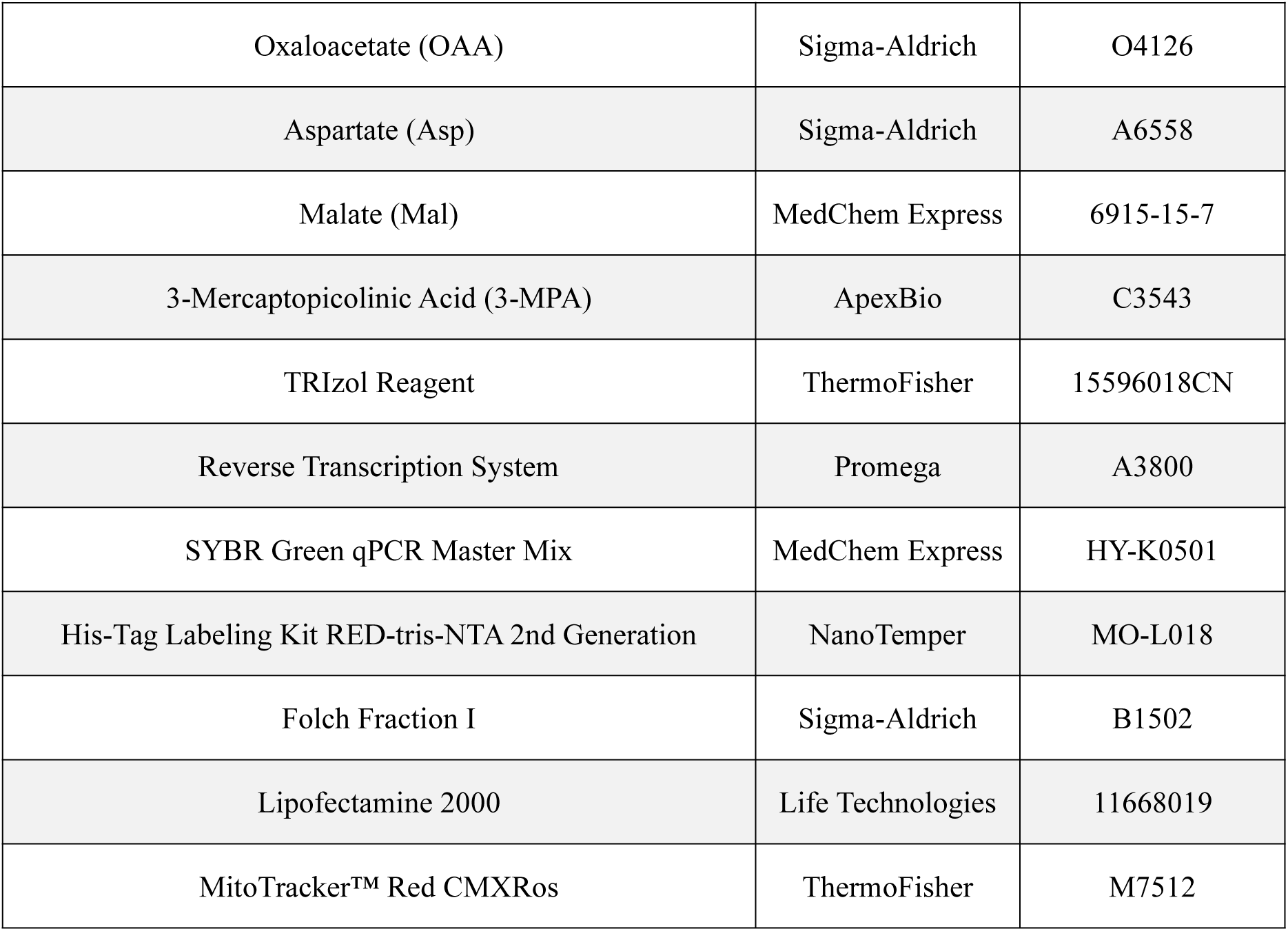
Antibodies and reagents used in this study.

| Antibodies | Antigen Species | Sources | Identifier |
| --- | --- | --- | --- |
| Mouse monoclonal anti-6×His-tag | Species independent | proteintech | 66005-1-Ig |
| Mouse monoclonal anti-GST-tag | Species independent | proteintech | 66001-2-Ig |
| Mouse monoclonal anti-HSP-60 | *C. elegans* | DHSB | AB_10572252 |
| Mouse monoclonal anti-α-Tubulin | Mouse, Human, Rat, Chicken, Bovine, Yeast and *C. elegans* | Sigma-Aldrich | T6199 |
| Mouse monoclonal anti-β-Actin | Mouse, Rat, Human, Drosophila, Yeast and *C. elegans* | Santa Cruz | sc-517582 |
| Mouse monoclonal anti-IMMT-1 | *C. elegans*, IMMT-1 (100-500 aa) | Homemade | This lab |
| Mouse monoclonal anti-CHCH-3 | *C. elegans*, Full length CHCH-3 | Homemade | This lab |
| Mouse monoclonal anti-TOM70 | Human, Mouse, Rat, Pig | proteintech | 66593-1-Ig |
| Goat Anti-Mouse IgG H&L (HRP) | Mouse | Abcam | ab205719 |
| Reagents |  | Sources | Identifier |
| ATP Assay Kit |  | Beyotime | S0026 |
| Carbonyl cyanide 4-(trifluoromethoxy)phenylhydrazone (FCCP) |  | MedChem Express | HY-100410 |
| Agilent Seahorse XF Calibrant (pH 7.4) |  | Agilent Technologies | 100840-000 |
| IPTG (Isopropyl β-D-thiogalactoside) | | MedChem Express | 367-93-1 |
| Imidazole |  | Sigma-Aldrich | 56750 |
| cOmplete™, EDTA-free Protease Inhibitor Cocktail |  | Roche | 4693132001 |
| Chelating Sepharose™ Fast Flow |  | GE Healthcare | 17-0575-01 |
| Glutathione Sepharose™ 4B |  | GE Healthcare | 17-0756-01 |
| GFP-Nanoab-Agarose Beads |  | Lablead | GNA-50-1000 |
| Oxaloacetate Assay Kit |  | Sigma-Aldrich | MAK070 |

|  |  |  |
| --- | --- | --- |
| Oxaloacetate (OAA) | Sigma-Aldrich | O4126 |
| Aspartate (Asp) | Sigma-Aldrich | A6558 |
| Malate (Mal) | MedChem Express | 6915-15-7 |
| 3-Mercaptopicolinic Acid (3-MPA) | ApexBio | C3543 |
| TRIzol Reagent | ThermoFisher | 15596018CN |
| Reverse Transcription System | Promega | A3800 |
| SYBR Green qPCR Master Mix | MedChem Express | HY-K0501 |
| His-Tag Labeling Kit RED-tris-NTA 2nd Generation | NanoTemper | MO-L018 |
| Folch Fraction I | Sigma-Aldrich | B1502 |
| Lipofectamine 2000 | Life Technologies | 11668019 |
| MitoTracker™ Red CMXRos | ThermoFisher | M7512 |

**S4 Table.**
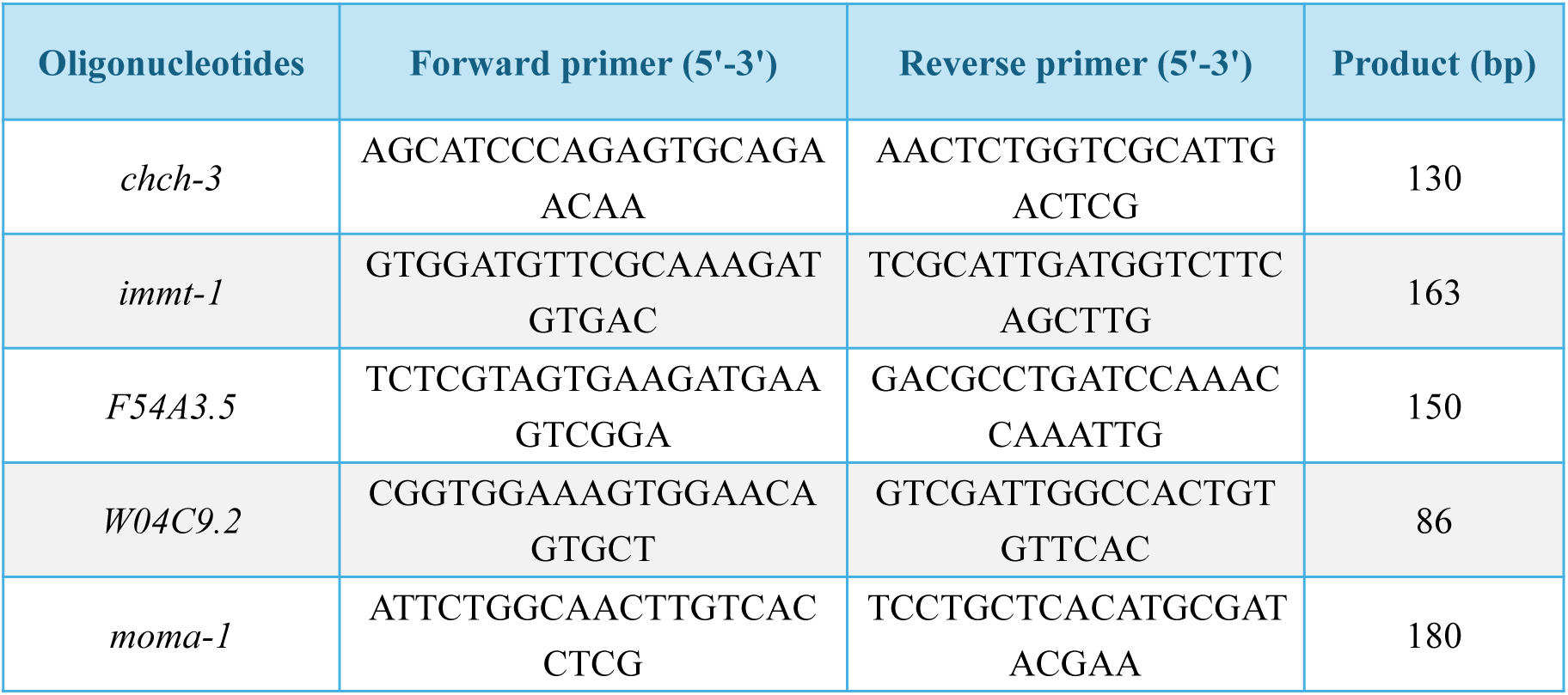
Oligonucleotides used for qRT-PCR in this study.

## Notes

### Competing Interest Statement

The authors have declared no competing interest.

